# Population genomics reveals complex patterns of immune gene evolution in monarch butterflies (*Danaus plexippus*)

**DOI:** 10.1101/620013

**Authors:** Wen-Hao Tan, Andrew J. Mongue, Jacobus C. de Roode, Nicole M. Gerardo, James R. Walters

**Affiliations:** Department of Biology, Emory University, Atlanta, Georgia, United States of America; Department of Ecology and Evolutionary Biology, University of Kansas, Lawrence, Kansas, United States of America

**Keywords:** immunity, natural selection, Lepidoptera, *Danaus*, ecological immunology

## Abstract

Immune genes presumably rapidly evolve as pathogens exert strong selection pressures on host defense, but the evolution of immune genes is also constrained by trade-offs with other biological functions and shaped by the environmental context. Thus, immune genes may exhibit complex evolutionary patterns, particularly when organisms disperse to or live in variable environments. We examined the evolutionary patterns of the full set of known canonical immune genes within and among populations of monarch butterflies (*Danaus plexippus*), and relative to a closely related species (*D. gilippus*). Monarchs represent a system with a known evolutionary history, in which North American monarchs dispersed to form novel populations across the world, providing an opportunity to explore the evolution of immunity in the light of population expansion into novel environments. By analyzing a whole-genome resequencing dataset across populations, we found that immune genes as a whole do not exhibit consistent patterns of selection, differentiation, or genetic variation, but that patterns are specific to functional classes. Species comparisons between *D. plexippus* and *D. gilippus* and analyses of monarch populations both revealed consistently low levels of genetic variation in signaling genes, suggesting conservation of these genes over evolutionary time. Modulation genes showed the opposite pattern, with signatures of relaxed selection across populations. In contrast, recognition and effector genes exhibited less consistent patterns. When focusing on genes with exceptionally strong signatures of selection or differentiation, we also found population-specific patterns, consistent with the hypothesis that monarch populations do not face uniform selection pressures with respect to immune function.

## 1 INTRODUCTION

The cellular and humoral immune systems provide one of the primary animal defenses against pathogens. Given that pathogens exert strong selection pressure on their hosts, immunity-related genes are presumed to be under selection and rapidly evolving due to host-pathogen coevolutionary arms races (McTaggart, Obbard, Conlon, & Little, 2012; Schlenke & Begun, 2003). However, the evolution of immune genes is also constrained by trade-offs with other biological functions and shaped by environmental context (Demas & Nelson, 2012). When animals colonize novel environments, they often encounter novel ecological conditions, including resources and pathogens, that could influence disease susceptibility and alter selection pressures on immune functions (Eizaguirre, Lenz, Kalbe, & Milinski, 2012). In addition to cellular and humoral immune defenses, animals may use behavioral defenses, medicinal compounds, and symbionts to protect against pathogens (Parker, Barribeau, Laughton, de Roode, & Gerardo, 2011). Utilization of alternative defenses may vary across populations due to environmental context, selection, plasticity, and genetic drift. These differences, in turn, could shape immune gene evolution across populations. Taken together, the evolutionary patterns of immune genes may be complicated, particularly when organisms disperse to novel environments.

The cellular and humoral immune system of insects is relatively simple compared to the vertebrate immune system, potentially facilitating study of immune gene evolution. The canonical immune system of insects mainly consists of four functional classes: recognition (e.g., peptidoglycan recognition proteins or PGRPs), signaling (e.g., the Toll signaling pathway), modulation (e.g., CLIP serine proteases), and effector (e.g., antimicrobial peptides: AMPs) (Christophides et al., 2002). Insect immune responses usually begin with the identification of foreign molecules by pattern recognition receptors encoded by recognition genes. The recognition of foreign molecules activates downstream signaling cascades that involve proteins encoded by signaling and modulation genes. For instance, recognition of Gram-positive bacteria and fungi often triggers the activation of the Toll signaling pathway, while recognition of Gram-negative bacteria often triggers the activation of the *immune deficiency* (IMD) signaling pathway. These signaling cascades lead to production of effector proteins (e.g., AMPs, pro-phenoloxidases that lead to melanization responses) that directly interact with pathogens (Lemaitre & Hoffmann, 2007).

Some studies of insect immune gene evolution have demonstrated that immune genes rapidly evolve. For example, Erler et al. (2014) showed that AMPs evolve much faster than non-immune genes in multiple bumblebee species, and Viljakainen et al. (2009) demonstrated that a select subset of immune genes (14 recognition and effector genes) are rapidly evolving in both honey bees and ants. However, these studies and most others have focused on only a few genes or one part of the immune system, without consideration of the full set of canonical immune genes.

Consideration of the immune gene set as a whole is important, in part, because different immune components may face different selection pressures. Specifically, coevolutionary theory would predict that molecules that directly interact with rapidly evolving pathogens – such as those encoded by recognition and effector genes – may undergo faster evolution than those involved in signal transduction. Indeed, a comparative study of twelve *Drosophila* species found that recognition proteins and effectors are rapidly evolving and highly differentiated; in contrast, proteins within signaling transduction cascades are more constrained across species (Sackton et al., 2007).

To our knowledge, only a few studies have taken a comprehensive, population-centered approach: Early et al. (2017) and Keehnen et al. (2018) examined the evolution of the full set of canonical immune genes across populations in fruit flies (*Drosophila melanogaster*) and a butterfly (*Pieris napi*), respectively. Studies on both species demonstrated that immune gene functional classes vary in their patterns of selection and differentiation, with conservation of signaling genes, balancing selection acting on effector genes, and recognition genes showing higher levels of between-population differentiation (Chapman, Hill, & Unckless, 2018; Early et al., 2017; Keehnen et al., 2018; Unckless, Howick, & Lazzaro, 2016).

In this study, we examined evolution of the full set of canonical immune genes across natural populations of monarch butterflies (*Danaus plexippus*). Monarchs are widely distributed, specialist herbivores that feed on toxic milkweed plants during their larval stage (Ackery & Vane-Wright, 1984; Oberhauser & Solensky, 2004). Monarchs originated in North America and colonized worldwide locations in the 19^th^ century through independent dispersal events across the Pacific Ocean, the Atlantic Ocean, and Central-South America (Fig. 1) (Ackery & Vane-Wright, 1984; Zhan et al., 2014), providing an opportunity to study immune gene evolution in the context of a known evolutionary history. Importantly, through these dispersal events, monarchs formed populations in which they relied on more toxic milkweed host plants and in which they experienced greater risk of infection by the common monarch parasite *Ophryocystis elektroscirrha* (Altizer & de Roode, 2015), likely altering selection on the monarch immune system. Here, we assessed patterns of divergence, diversity, and selection for monarch immune genes, using *D. gilippus* as an outgroup and contrasting the ancestral North American monarch population with geographically and genetically distinct derived populations.

**Figure 1.**
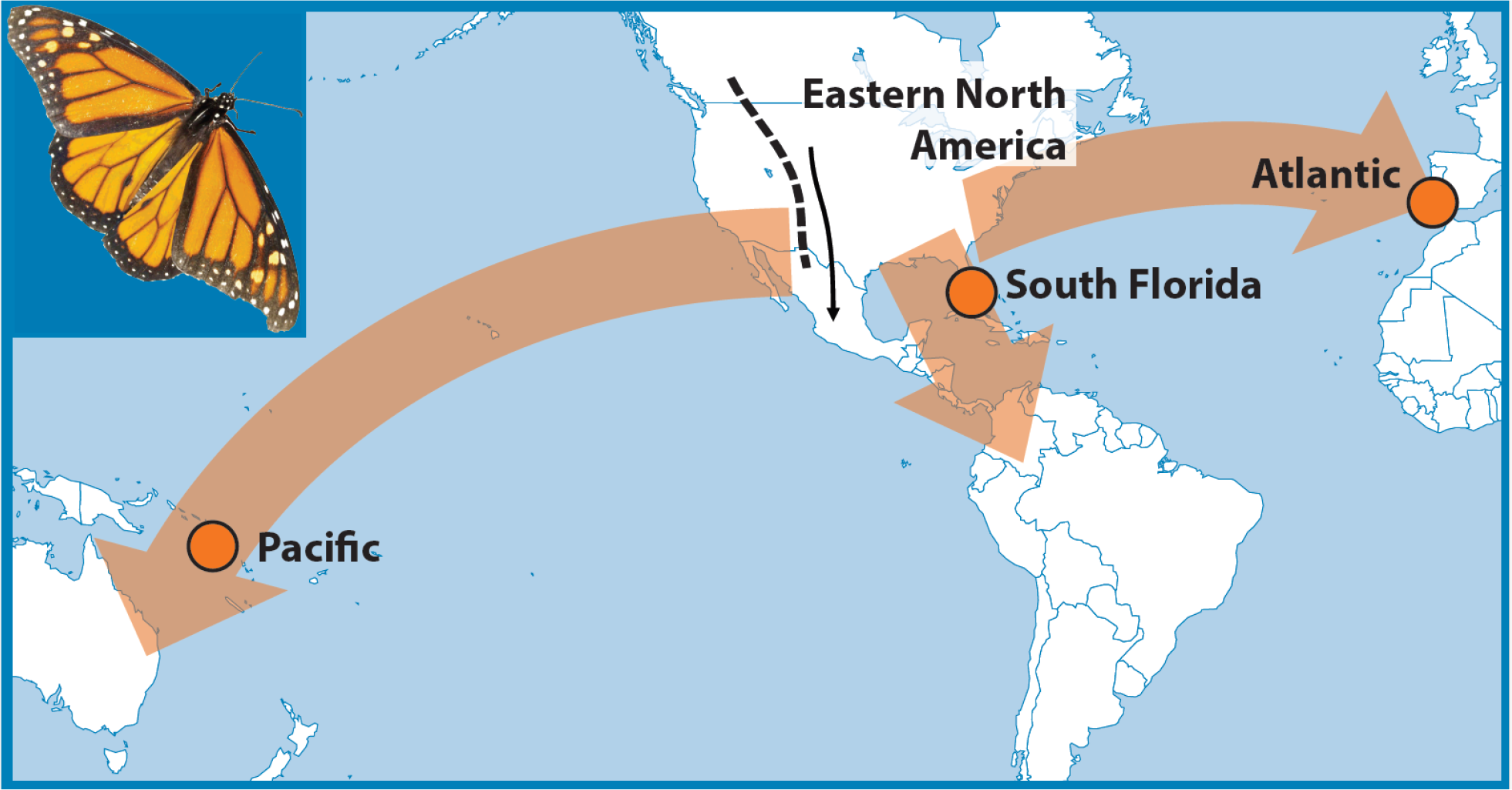
Current distribution of monarch butterflies around the world and their historical dispersal routes. Monarchs originated in the North America and established other populations via three main dispersal events: across the Pacific Ocean, across the Atlantic Ocean, and toward Central/South America.

## 2 MATERIALS AND METHODS

### 2.1 Overview of approach

Differential selection pressures owing to ecological differences could affect the type and strength of selection on immune genes. In addition to selection, other factors such as demographic history and local genomic factors also may affect their evolutionary patterns. Given that several population genetic measures of selection are sensitive to demographic effects, past demographic history and recent dispersal are important factors that could influence and/or confound observed signatures of selection (Eyre-Walker & Keightley, 2009; Vitti, Grossman, & Sabeti, 2013). In most population genomic studies of immune genes, relatively little is known about the demographic history and the ecological differences of the focal populations; however, in monarch butterflies, previous population genetic and genomic studies have inferred that monarchs originated in North America and recently spread around the world via three major dispersal events (Pierce et al., 2014; Zhan et al., 2014). While these events led to formation of populations subject to different ecological conditions, the dispersal process itself may also influence patterns of population genetics. To account for this, we used a paired-control approach to determine if signatures of selection in functional classes of immune genes differ from those in the background genome. In addition, we identified individual immune genes that are genome-wide outliers for combinations of population genetic parameters, indicating they are likely experiencing different selective pressures.

### 2.2 The population genomic dataset

We obtained a whole genome Illumina sequencing dataset from Zhan et al. (2014), who sequenced monarch samples across populations worldwide. Based on previous population genetic and genomic studies (Pierce et al., 2014; Zhan et al., 2014), we assigned monarch samples into genetic populations according to their collection location. We excluded samples with average sequencing depth lower than 10X for quality control purposes. We used a total of 37 whole monarch genomes in our study, including the ancestral population (North America) and derived populations in South Florida, the Pacific, and the Atlantic (Fig. 1, supplemental information Table S1).

We aligned sequencing reads to the reference monarch genome (Zhan, Merlin, Boore, & Reppert, 2011) using Bowtie2 with the option “--very-sensitive-local” (Langmead & Salzberg, 2012). After reference mapping, we took the alignments through the Genome Analysis Tool Kit’s best practices pipeline to remove PCR duplicates and realign around variable insertions and deletions (McKenna et al., 2010).

### 2.3 Gene sets

We obtained a full set of annotated monarch immune genes published by the *Heliconius* Genome Consortium (2012), which included a set of annotated (*Heliconius*) immune genes and their orthologs in several species, including monarchs. The monarch orthologs listed in this published dataset were based on a previous version of monarch genome annotation (OGS1.0), so we updated this full set of immune genes to the latest version of gene annotation (OGS2.0) using information provided in Monarch Base (Zhan & Reppert, 2013). This updated monarch immune gene set contains 114 genes belonging to functional classes of recognition, signaling, modulation, and effector (see supplemental information Table S2). We also obtained the latest version (OGS2.0) of all the annotated monarch genes from the published reference genome (Zhan et al., 2011; Zhan & Reppert, 2013) in order to compare evolution of immune genes to evolution of non-immune genes (as controls) in the background genome.

We restricted our analyses to autosomal genes to avoid the complication of unequal sampling between autosomes and the Z sex chromosome; sequenced individuals were of different sexes, so a variable number of Z chromosomes were sampled. We did not perform a separate analysis of Z-linked genes due to sample size limitations. We identified Z-linked immune genes based on chromosomal assignments obtained from Mongue et al. (2017). The majority of immune genes are on autosomes, with only 12 genes located on the Z chromosome (see supplemental information Table S2).

### 2.4 Population genetic analyses

We calculated four population genetic statistics: pairwise nucleotide diversity (π), Watterson’s θ (Nei, 1979; Watterson, 1975), Tajima’s D (Tajima, 1989), and *F*_ST_ (Wright, 1921). We generated folded site frequency spectra (SFS) and calculated the four statistics using ANGSD (Korneliussen, Albrechtsen & Nielsen, 2014). We calculated π, Watterson’s θ and Tajima’s D for each population; we calculated *F*_ST_ between populations by comparing each of the three derived populations (*i.e.*, Florida, Pacific, and Atlantic) to the ancestral population (North America). For all calculations, we first generated a SFS for all genes in the same functional class to use as a prior for gene-specific parameter estimates. Using this prior, we then calculated those four population genetic statistics for each gene in the functional class. We repeated the procedures for each gene with either: (1) 0-fold degenerate sites; (2) 4-fold degenerate sites; and (3) all sites within each gene. The 0-fold and 4-fold degeneracy sites for all monarch genes were obtained from Mongue et al. (2019). The genomic position of each gene was obtained from the latest version of gene annotation (OGS2) in Monarch Base (Zhan et al., 2011). We performed all calculations for all genes in the genome. We generated inputs for ANGSD and processed the data using custom R and python scripts in R version 3.4.1 (R Core Team, 2017) and python version 2.7.5.

### 2.5 A paired control approach to compare immune genes to the genomic background

Evolutionary change of a gene can be influenced by gene length and local genomic factors, such as recombination rate and selection on nearby genes (Castellano, Coronado-Zamora, Campos, Barbadilla, & Eyre-Walker, 2016; Comeron, Ratnappan, & Bailin, 2012; Wong et al., 2008). Therefore, we evaluated whether immune genes differed from broader patterns in the genome background using a paired-control approach that compares immune genes to a selected subset of control genes. This paired-control approach enables us to take these factors into consideration, assessing the patterns of selection more conservatively; our approach is similar to that used by Early et al. (2017) and Chapman, Hill, & Unckless (2018).

Specifically, we first constructed a pool of control genes for each immune gene based on the following criteria: (1) the length of the control genes are within either 0.5-2 times, or ±1500 bp, of the total length of the immune gene; (2) control genes are on the same scaffold (and thus chromosome) as the immune gene; (3) control genes are not known to have immune function. Given that a high proportion of scaffolds in the reference monarch genome are relatively small in size (N50 = 715 kbp) (Zhan & Reppert, 2013), in some cases control gene pools were small. When a candidate gene pool was smaller than eight genes, we relaxed the location criterion and expanded the search to the whole chromosome level, while keeping the other two criteria unchanged. In all cases, we were able to gather > 8 candidate genes. Four focal immune genes did not have a chromosomal assignment. For these, we searched for genes that also did not have chromosomal assignments that fit the size and gene function criteria. We excluded genes that did not have an adequate number of 0-fold or 4-fold sites for estimating population genetic statistics from the control gene pools. For a given immune functional group, we calculated the test statistic as the summation of the difference between an immune gene and the mean of its control genes. We determined significance through 10,000 permutations. For each permutation round, we randomly sampled one gene for each immune gene from a pool containing the immune gene itself and all corresponding control genes with replacement to serve as the test gene, and calculate the difference between the test gene and the mean of the remaining genes in the pool. The permuted test statistic is calculated as the summation of those differences for genes belonging to a given immune functional group. We calculated *P*-values as the percentage of the 10,000 permutations in which the absolute value of the test statistic (observed value) is less than the absolute mean value of the permuted sets (permuted null distribution). The paired-control analyses were performed using custom R scripts in R version 3.4.1 (R Core Team, 2017).

### 2.6 Between-species analyses

In addition to between-population comparisons, we also sought to estimate longer-term evolutionary patterns by leveraging whole genome sequencing of a congener, the queen butterfly (*Danaus gilippus*) (Zhan et al., 2014). This gave us the opportunity to look at scaled rates of divergence between species (Dn/Ds). We aligned *D. gilippus* reads to the monarch reference using the stampy alignment software (Lunter & Goodson, 2011), parameterized for an increased (10%) substitution rate between reads and reference. These data were then taken through GATK’s best practice pipeline for SNP calling, including quality filtering of variants (McKenna et al., 2010). Passing variants were classified as synonymous or non-synonymous by SNPeff (Cingolani et al., 2012). Finally, we calculated Dn per gene as the number of nonsynonymous substitutions per non-synonymous site (and likewise for Ds), using previous knowledge of the degeneracy of each position in a coding sequence (Mongue et al., 2019). Owing to a substantial number of genes in both the immune and control sets with undefined Dn/Ds (created by zero counts of Ds), we did not implement a paired permutation test. Rather, we used R to perform Mann-Whitney U tests to assess significance of differences in divergence rates for immune gene classes compared to the control genes with non-zero synonymous divergence.

The *D. gilippus* data additionally allowed us to estimate the proportion of substitutions driven by adaptation (α) for immune genes and to compare with estimates from corresponding control genes in the monarch genome. As with established methods (Mongue et al., 2019), we used the queen butterfly sequences to infer a parsimonious ancestral (allele) state at polymorphic sites in the monarch genome, allowing us to generate an unfolded SFS, *i.e.* one that differentiates ancestral and derived allele frequencies. With unfolded spectra, we employed the likelihood model implemented in polyDFE (Tataru, Mollion, Glémin, & Bataillon, 2017) to estimate α and the distribution of fitness effects of new non-synonymous mutations (DFEs) while accounting for demography and errors in allele frequency polarization. To assess uncertainty in these estimates, parametric bootstrapping of input SFS (as implemented in polyDFE) was used to obtain a distribution of α and DFE statistics. Significant differences in α were apparent based on the non-overlapping confidence intervals of immune and control sets and did not warrant further statistical testing. Differences between DFEs were not formally tested but were used as ancillary, qualitative inferences to contextualize related results. Bootstrapping, statistical analyses, and visualization were completed with custom R scripts.

### 2.7 Outlier analyses

To identify specific loci that may experience distinctive evolutionary pressures, we searched for immune genes which are outliers relative to the genome-wide distributions of population genetic parameters. We jointly considered Tajima’s D and *F*_ST_, reasoning that loci showing extreme values for both parameters are likely to be of particular interest. We performed the analyses across all genes in the genome at either 0-fold sites or 4-fold sites and used information at genome-wide 0-fold or 4-fold sites as prior for estimating SFS in ANGSD. We converted Tajima’s D and *F*_ST_ values into percentiles in their genome-wide distribution. We defined genes that were either in the < 2.5^th^ percentile or >97.5^th^ percentile as genome outliers. To assess the outlier patterns considering both selection and population differentiation, we evaluated the relationship of Tajima’s D and *F*_ST_ for each functional class. Converting the values into percentiles also enabled us to compare patterns across populations. We visualized the patterns by plotting the Tajima’s D and *F*_ST_ genome percentiles against each other in a 2-D plot with Tajima’s D on the x-axis and *F*_ST_ on the y-axis. Separating the plot by the genome median of the two measures, it contains four quadrants: top-right (x > 0.5 & y > 0.5), bottom-right (x > 0.5 & y < 0.5), top-left (x < 0.5 and y > 0.5), and bottom-left (x < 0.5 and y < 0.5). Outliers falling into each of the four quadrants suggest different evolutionary scenarios: “top-right” suggests balancing selection acting differently between populations, “bottom-right” suggests balancing selection acting similarly between populations, “top-left” suggests directional selection acting differently between populations, and “bottom-left” suggests directional selection acting similarly between populations. We summarized the number of outliers in each area in contingency tables and analyzed the patterns. Due to small count numbers in some cells, we used Fisher’s exact tests. In addition, we examined whether immune genes are disproportionally represented in genome-wide outliers using Chi-square tests. All statistical analyses were performed in R.

## 3 RESULTS

### 3.1 North America: the ancestral population

#### A. Within-species analyses: characterizing genetic diversity and signatures of selection

As a group, immune genes showed slightly lower genetic diversity compared to paired-control genes, though this result was not statistically significant (Table 1 and Fig. 2). However, levels of genetic variation varied notably among the different functional classes of immune genes. At 0-fold sites, recognition and modulation genes exhibited a trend toward higher genetic variation than their respective control genes, while signaling and effector genes showed a trend toward lower genetic variation than their respective control genes. Signaling genes had a significantly lower π and Watterson’s θ at the 0-fold sites than controls, while other functional groups did not differ significantly from their controls. At 4-fold sites, none of the functional groups differed significantly from their controls; only the signaling genes had a marginally significantly lower π compared to controls.

**Table 1.** Population genetic statistics of immune genes in the North American population using the paired-control approach. The upper half shows results based on the 0-fold sites and the lower half shows results based on the 4-fold sites. The *F*_ST_ section is non-applicable because the North American population was the reference population used for population comparisons. “All immune” indicates the full immune gene set. In each statistic, the first row shows the test statistic of the immune gene group. The second row shows the proportion of 10,000 permutations in which the difference between the means of the immune gene group and the control set was positive. Percentages < 2.5% and > 97.5 % are labeled in bold. The third row shows the *P*-value. *P*-values < 0.05 are labeled in bold. Asterisks indicate: * < 0.05, ** < 0.01, *** < 0.001.

|  | All Immune | Recognition | Signaling | Modulation | Effector |
| --- | --- | --- | --- | --- | --- |
| 0-fold sites |  |  |  |  |  |
| $\pi$: test statistic | -0.07 | 0.01 | -0.07 | 0.01 | -0.02 |
| $\pi$: > 0 (%) | 5.84 | 83.10 | **0.04** | 75.68 | 14.60 |
| $\pi$: $P$-value. | 0.140 | 0.340 | **0.024\*** | 0.508 | 0.311 |
| Watterson's $\theta$: test statistic | -0.09 | 0.02 | -0.09 | 0.02 | -0.04 |
| Watterson's $\theta$: > 0 (%) | 3.71 | 88.83 | **0.03** | 76.02 | 6.53 |
| Watterson's $\theta$: $P$-value | 0.092 | 0.184 | **0.012\*** | 0.497 | 0.168 |
| Tajima's D: test statistic | -4.25 | -8.01 | -2.38 | 0.69 | 5.44 |
| Tajima's D: > 0 (%) | 30.29 | **0.76** | 33.12 | 58.16 | 94.66 |
| Tajima's D: $P$-value | 0.599 | **0.024\*** | 0.643 | 0.859 | 0.098 |
| $F_{ST}$: test statistic | NA | NA | NA | NA | NA |
| $F_{ST}$: > 0 (%) | NA | NA | NA | NA | NA |
| $F_{ST}$: $P$-value | NA | NA | NA | NA | NA |
| 4-fold sites |  |  |  |  |  |
| $\pi$: test statistic | -0.11 | -0.04 | -0.14 | 0.03 | 0.03 |
| $\pi$: > 0 (%) | 16.18 | 21.16 | 2.55 | 72.58 | 71.43 |
| $\pi$: $P$-value. | 0.327 | 0.419 | 0.059 | 0.564 | 0.596 |
| Watterson's $\theta$: test statistic | -0.09 | -0.01 | -0.10 | 0.06 | -0.04 |
| Watterson's $\theta$: > 0 (%) | 23.48 | 44.04 | 9.57 | 80.66 | 25.65 |
| Watterson's $\theta$: $P$-value | 0.467 | 0.856 | 0.192 | 0.387 | 0.510 |
| Tajima's D: test statistic | -8.60 | -4.41 | -10.33 | -1.07 | 7.21 |
| Tajima's D: > 0 (%) | 13.10 | 9.24 | **1.87** | 38.65 | **98.87** |
| Tajima's D: $P$-value | 0.262 | 0.189 | **0.043\*** | 0.760 | **0.015\*** |
| $F_{ST}$: test statistic | NA | NA | NA | NA | NA |
| $F_{ST}$: > 0 (%) | NA | NA | NA | NA | NA |
| $F_{ST}$: $P$-value | NA | NA | NA | NA | NA |

**Figure 2.**
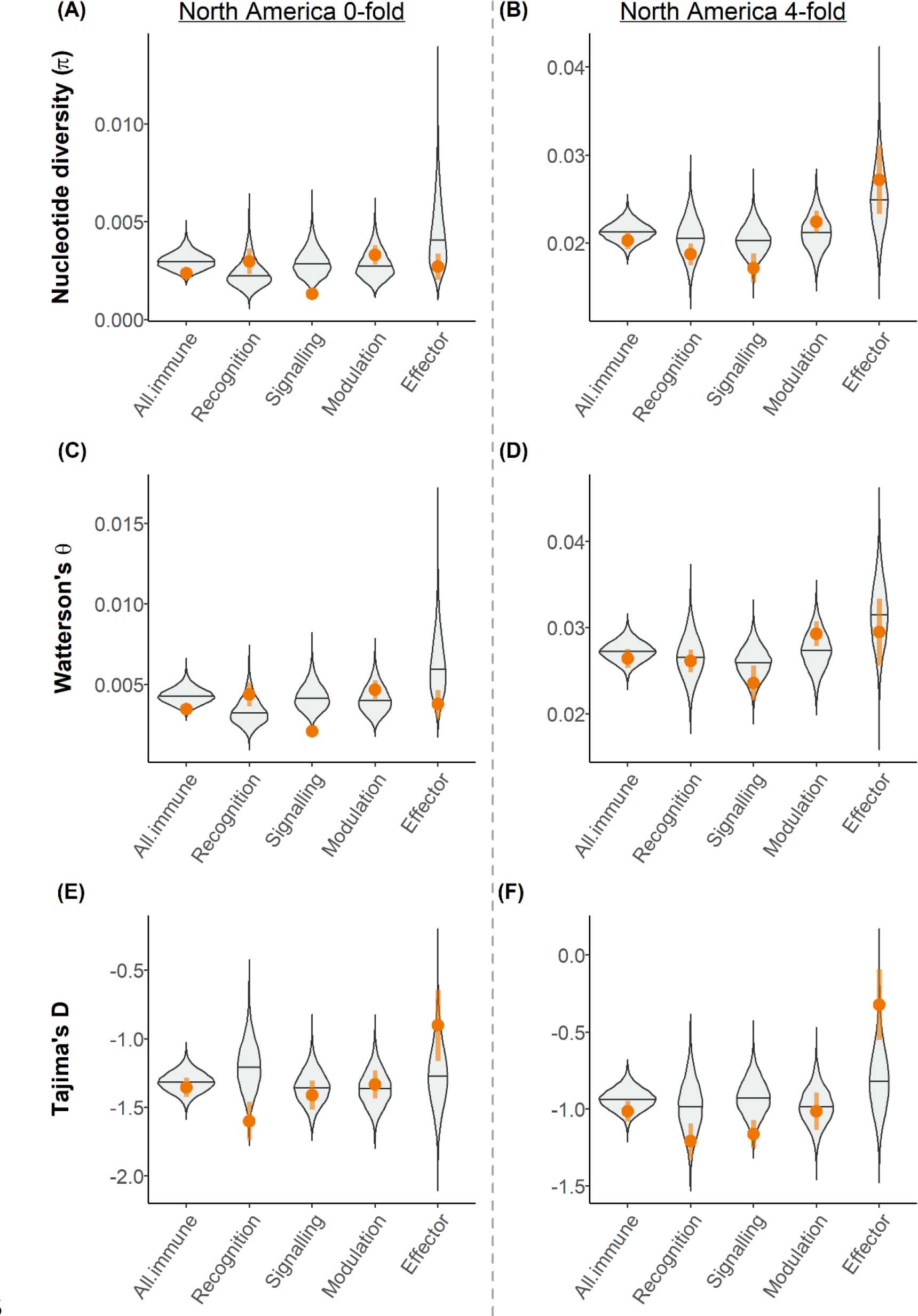
Population genetic statistics of immune genes in the North American population using the paired-control approach. 0-fold sites shown in (A), (C), (E) and 4-fold sites shown in (B), (D), (F). (A)-(B): Nucleotide diversity (**π**); (C)-(D): Watterson’s θ; (E)-(F): Tajima’s D. Each immune gene group was compared to selected pair-control sets. Violin plots show the distribution of the mean of each control set generated with 10, 000 permutations. The orange dots and vertical lines indicate mean ±1 SEM of the immune gene group of interest.

Immune genes as a whole did not show a distinct pattern of selection; the full set of immune genes was not significantly different from the paired-controls (Table 1 and Fig. 2). However, as with π and Watterson’s θ, patterns of Tajima’s D varied across different functional classes of immune genes. Recognition genes showed a trend of lower Tajima’s D at both 0-fold and 4-fold sites but was only significantly lower at the 0-fold sites. Signaling genes showed a significantly lower Tajima’s D than controls at only the 4-fold sites. Modulation genes did not exhibit any significant differences to the controls. Effector genes showed significantly higher Tajima’s D at the 4-fold sites and marginally significantly higher Tajima’s D at the 0-fold sites to their respective controls.

Taken together, the full set of immune genes did not differ from control genes in either genetic diversity or signatures of selection; however, different functional classes exhibited significant differences. Specifically, signaling genes showed lower genetic variation than control genes, consistent with broad purifying selection; associated background selection could explain the reduced 4-fold site Tajima’s D. By contrast, the strongly elevated Tajima’s D among effector genes seems best explained by frequent balancing selection among these loci. Analyses based on all sites within each gene showed similar qualitative results (see supplemental information Table S3 and Fig. S1).

#### B. Between-species analyses: comparing D. plexippus and D. gilippus

We further assessed molecular evolutionary patterns of immune genes by estimating divergence to the closely related queen butterfly. We tested for differences in rates of divergence (Dn/Ds) between immune genes and their controls selected from the rest of the genome. We found that neither effector (Fig. 3; W = 2866, *P* = 0.764) nor signaling genes (Fig. 3; W = 23427, *P* = 0.352) showed increased divergence compared to their controls, which is consistent with balancing and purifying selection respectively decreasing the fixation rate of variants. In contrast, we found elevated divergence in both modulation (Fig. 3; W = 6036.5, *P* = 0.009) and recognition genes (Fig. 3; W = 1156, *P* = 0.018) compared to their controls. Such a result is indicative of either increased directional selection or relaxed constraint allowing more non-synonymous differences to reach fixation. Taken together with within-species analyses of nucleotide diversity, these results suggest that relaxed selection is more likely for modulation genes, but the cause of increased divergence in recognition genes is less immediately apparent.

**Figure 3.**
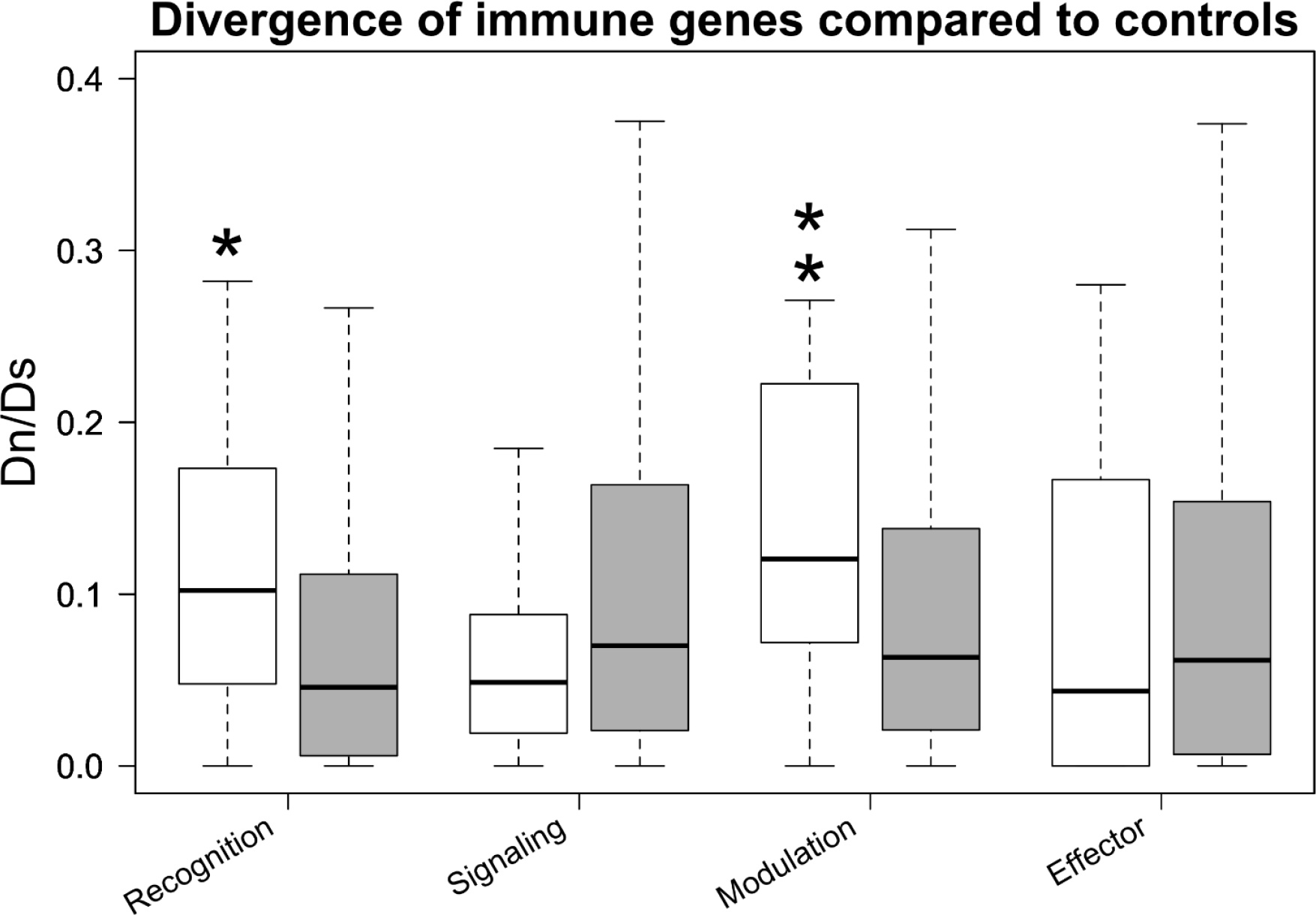
Divergence rates compared for immune genes and paired-controls using the queen butterfly as a reference. Here Dn is calculated as non-synonymous substitutions per non-synonymous site, and Ds is the number of synonymous substitutions per synonymous site. Rates for each gene class are labeled, with the control group in grey immediately to right. Asterisks represent levels of significance in a Mann-Whitney-U Test following the convention: * for <0.0.5 and ** < 0.01.

#### C. Distributions of fitness effects and estimates of adaptive evolution

To further investigate patterns of selection, we used SFS to estimate the distribution of fitness effects for new non-synonymous mutations (DFEs) among the immune gene functional classes and their control sets. Though we are unable to statistically compare differences between immune gene groups and their controls, the patterns are largely consistent with the results of other tests. Signaling genes exhibited a lack of neutral and weakly selected variants, combined with an increase in strongly deleterious and (to a lesser degree) beneficial variants (Fig. 4, second row). This pattern suggests most new variation is destined to be removed by purifying selection, with occasional adaptive fixations. Modulation genes did not greatly differ from their control set, though the slight increase in inferred neutral and weakly selected variants (−10 < s < 1) is consistent with relaxed selection in this class of genes (Fig. 4, third row). Effector genes, however, showed a lack of strongly deleterious (s < −100) and an increase in moderately deleterious (−100 < s < −10) variants (Fig. 4, fourth row). This dearth of strongly deleterious variants suggests that alleles can reach more intermediate frequency, as expected under balancing selection.

**Figure 4.**
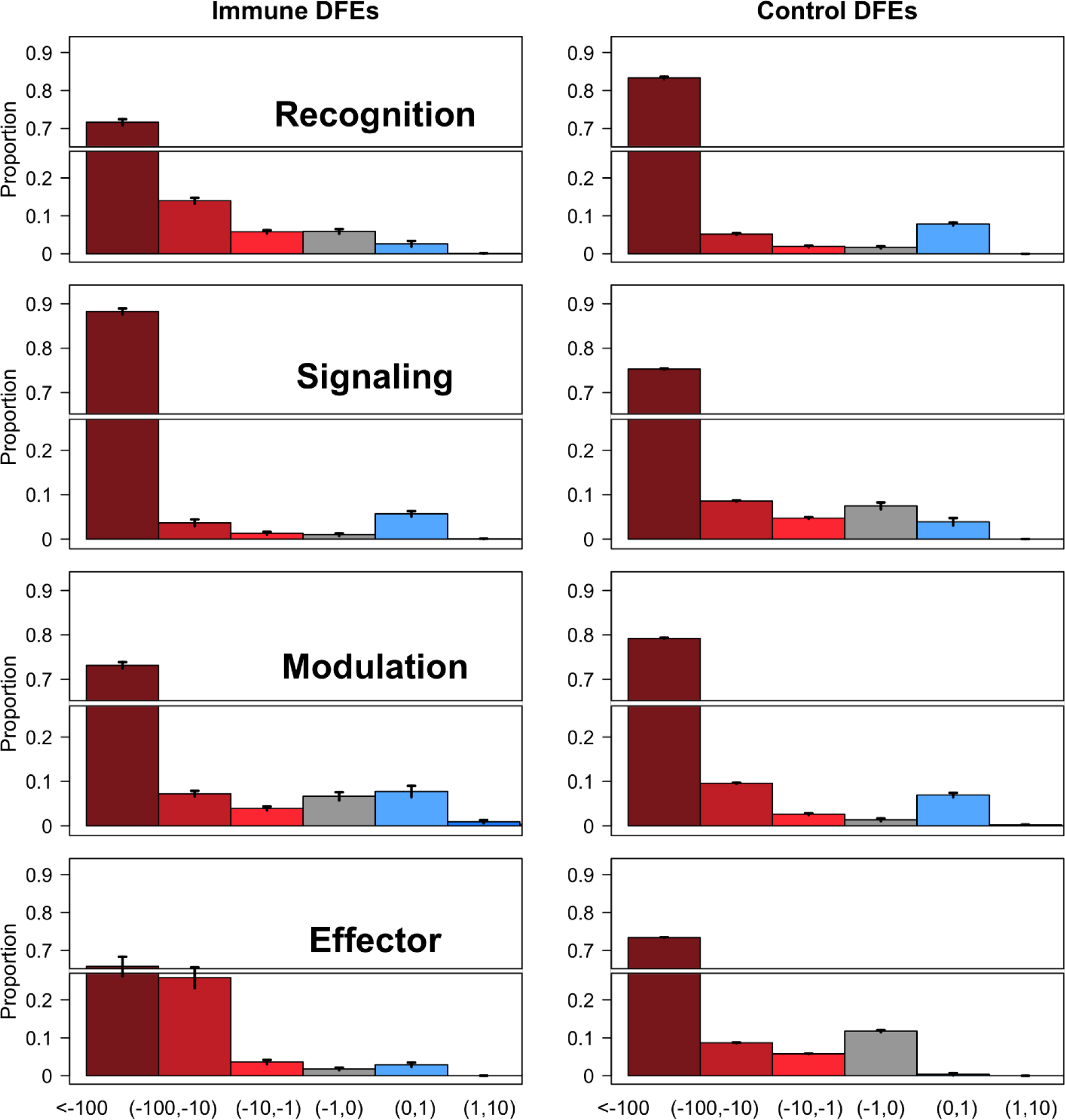
Predicted distributions of fitness effects of new, non-synonymous mutations for each of the four classes of immune genes (left column) and their paired-control sets (right column). Bars represent the proportion of variants that fall within a given selective class (s), from strongly deleterious (far left, darkest red) to beneficial (right, blue). Each plot is scaled with the same y-axis and has a gap from 0.25 to 0.65 to allow visualization of the whole distribution. Vertical lines on each bar, while mostly too small to notice, represent twice the standard error of the mean per-selective-class estimate from one hundred parametric bootstrap replicates. Estimates come from the tool polyDFE.

Unlike the other classes of immune genes, the DFEs for recognition genes and their controls suggest an alternative explanation for the patterns observed in other population genetic statistics. Note that here, the control genes (Fig. 4, top right) exhibit a similar pattern to the one described above for the focal set of signaling genes, *i.e.* purifying selection. The recognition genes’ DFEs, however, do not appear to be skewed by strong selection. In this light, other results for recognition genes may have more to do with purifying selection on controls than on selection on the recognition genes themselves.

Finally, we used the DFEs to estimate α (the proportion of adaptive substitutions) in each immune class and its control set. We found that α was significantly different between immune genes and controls in each of the four groups, as evidenced by non-overlapping confidence intervals (Fig. 5). For three of the four classes, the direction of these differences is consistent with other lines of evidence for selection. Namely, we found more adaptive evolution in effector and signaling genes and less adaptation in modulation genes compared to their controls. For recognition, however, evidence for less adaptation than controls conflicts with the evidence for selection from Tajima’s D. This lower α, alongside the DFEs, suggest that recognition genes are under weaker selection than their paired controls.

**Figure 5.**
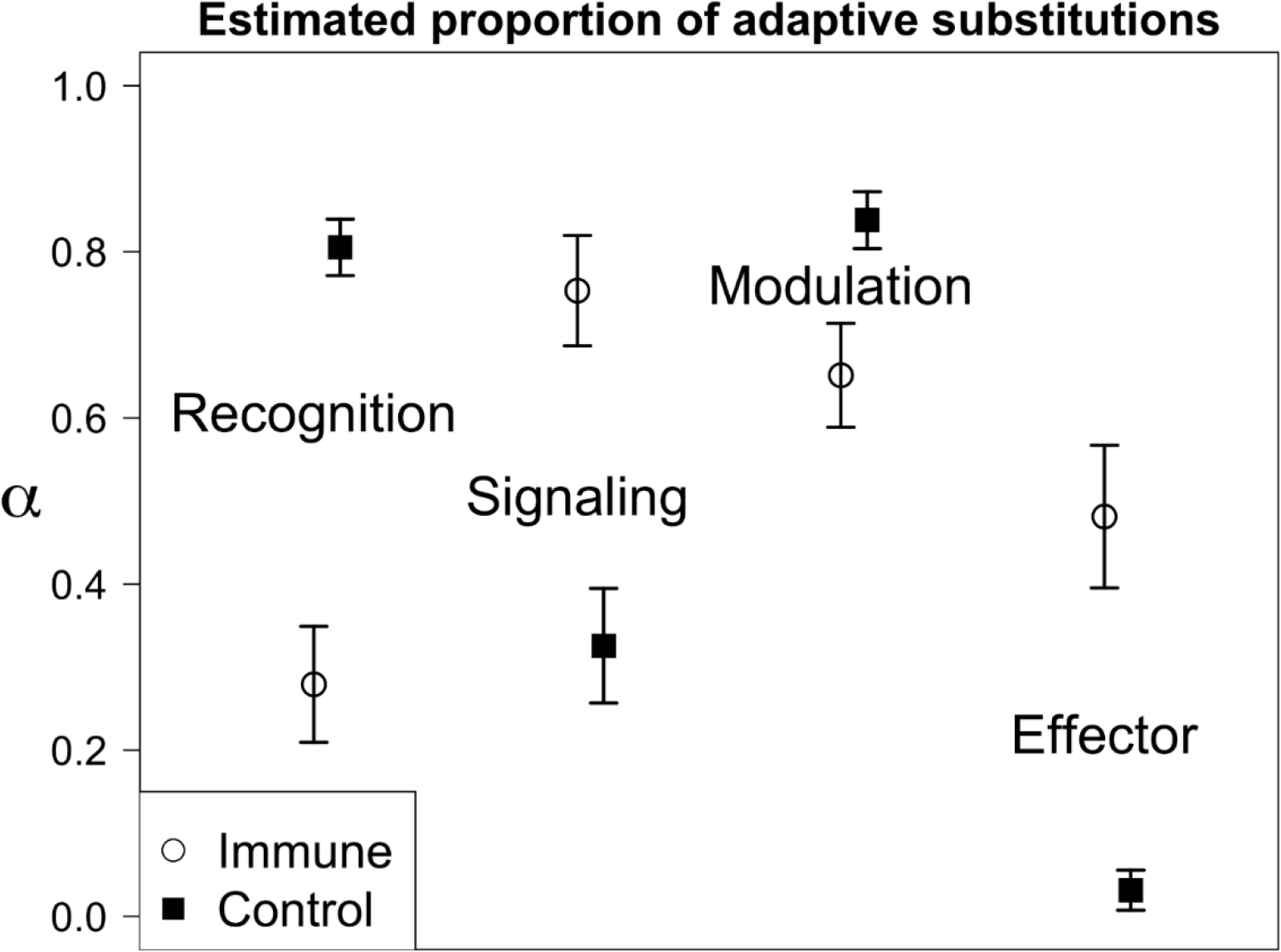
Estimates of the proportion of substitutions resulting from adaptive processes (α) based on DFEs computed in polyDFE. Each immune gene class (open circles) has a paired-control set of genes immediately to its right (filled squares). Error bars represent twice the standard error of the mean of one hundred parametric bootstrap replicates of the input data (site frequency spectra). All immune-control comparisons are significantly different from zero and each immune class is significantly different from its controls.

### 3.2. Population-level comparisons: the ancestral and three derived populations

#### A. Within-population analyses: characterizing genetic diversity and signatures of selection

Consistently across all four populations, the full set of immune genes did not show any significant differences compared to control genes at either 0-fold or 4-fold sites (Tables 1–4 and Fig. 6). For recognition genes, there was an overall trend toward higher genetic variation than controls at the 0-fold sites across populations; however, this was not statistically significant for any population. For signaling genes, there was an overall consistent trend toward lower genetic variation than controls at both the 0-fold and 4-fold sites across populations. Notably, in all populations, both π and Watterson’s θ were significantly lower than controls at the 0-fold sites of signaling genes. For modulation genes there was an overall trend toward higher genetic variation than controls across populations for both the 0-fold and 4-fold sites; however, this was not statistically significant for any of the four populations. For effector genes, the pattern was more variable, and no significant differences to the controls were found in any of the populations.

**Table 2.** Population genetic statistics of immune genes in the Florida population using the paired-control approach. The upper half shows results based on the 0-fold sites and the lower half shows results based on the 4-fold sites. *F*_ST_ was compared to the North American population. “All immune” indicates the full immune gene set. In each statistic, the first row shows the test statistic of the immune gene group. The second row shows the proportion of 10,000 permutations in which the difference between the means of the immune gene group and the control set was positive. Percentages < 2.5% and > 97.5 % are labeled in bold. The third row shows the *P*-value. *P*-values < 0.05 are labeled in bold. Asterisks indicate: * < 0.05, ** < 0.01, *** < 0.001.

|  | All Immune | Recognition | Signaling | Modulation | Effector |
| --- | --- | --- | --- | --- | --- |
| 0-fold sites |  |  |  |  |  |
| $\pi$: test statistic | -0.07 | 0.01 | -0.07 | 0.01 | -0.03 |
| $\pi$: > 0 (%) | 4.54 | 78.83 | **0.05** | 76.22 | 13.21 |
| $\pi$: $P$-value. | 0.118 | 0.461 | **0.025\*** | 0.498 | 0.291 |
| Watterson's $\theta$: test statistic | -0.08 | 0.02 | -0.08 | 0.01 | -0.03 |
| Watterson's $\theta$: > 0 (%) | 3.44 | 82.81 | **0.07** | 73.52 | 5.98 |
| Watterson's $\theta$: $P$-value | 0.090 | 0.346 | **0.021\*** | 0.558 | 0.161 |
| Tajima's D: test statistic | -5.02 | -6.80 | -11.83 | 1.14 | 12.46 |
| Tajima's D: > 0 (%) | 29.28 | 3.40 | 2.78 | 60.66 | **99.91** |
| Tajima's D: $P$-value | 0.594 | 0.077 | 0.062 | 0.794 | **0.001\*\*** |
| $F_{ST}$: test statistic | 0.59 | 0.53 | 0.14 | 0.03 | -0.11 |
| $F_{ST}$: > 0 (%) | 94.37 | **99.84** | 71.61 | 61.59 | 18.02 |
| $F_{ST}$: $P$-value | 0.098 | **0.002\*\*** | 0.575 | 0.837 | 0.349 |
| 4-fold sites |  |  |  |  |  |
| $\pi$: test statistic | -0.05 | -0.01 | -0.13 | 0.06 | 0.02 |
| $\pi$: > 0 (%) | 34.57 | 44.02 | 3.79 | 85.30 | 66.74 |
| $\pi$: $P$-value. | 0.683 | 0.868 | 0.090 | 0.294 | 0.706 |
| Watterson's $\theta$: test statistic | 0.00 | 0.02 | -0.10 | 0.09 | -0.02 |
| Watterson's $\theta$: > 0 (%) | 50.89 | 69.13 | 10.34 | 93.40 | 38.79 |
| Watterson's $\theta$: $P$-value | 0.979 | 0.629 | 0.209 | 0.129 | 0.764 |
| Tajima's D: test statistic | -9.01 | -4.80 | -7.70 | -2.76 | 6.26 |
| Tajima's D: > 0 (%) | 14.05 | 5.21 | 7.91 | 25.61 | 95.51 |
| Tajima's D: $P$-value | 0.281 | 0.107 | 0.165 | 0.501 | 0.083 |
| $F_{ST}$: test statistic | -0.22 | -0.12 | 0.05 | -0.14 | -0.02 |
| $F_{ST}$: > 0 (%) | 24.36 | 15.97 | 61.89 | 15.73 | 45.88 |
| $F_{ST}$: $P$-value | 0.470 | 0.321 | 0.810 | 0.314 | 0.849 |

**Table 3.** Population genetic statistics of immune genes in the Pacific population using the paired-control approach. The upper half shows results based on the 0-fold sites and the lower half shows results based on the 4-fold sites. *F*_ST_ was compared to the North American population. “All immune” indicates the full immune gene set. In each statistic, the first row shows the test statistic of the immune gene group. The second row shows the proportion of 10,000 permutations in which the difference between the means of the immune gene group and the control set was positive. Percentages < 2.5% and > 97.5 % are labeled in bold. The third row shows the *P*-value. *P*-values < 0.05 are labeled in bold. Asterisks indicate: * < 0.05, ** < 0.01, *** < 0.001.

|  | All Immune | Recognition | Signaling | Modulation | Effector |
| --- | --- | --- | --- | --- | --- |
| 0-fold sites |  |  |  |  |  |
| $\pi$: test statistic | -0.05 | 0.02 | -0.06 | 0.01 | -0.02 |
| $\pi$: > 0 (%) | 11.28 | 90.28 | **0.09** | 72.45 | 32.43 |
| $\pi$: $P$-value. | 0.239 | 0.139 | **0.032\*** | 0.598 | 0.546 |
| Watterson's $\theta$: test statistic | -0.04 | 0.02 | -0.05 | 0.01 | -0.01 |
| Watterson's $\theta$: > 0 (%) | 9.07 | 96.35 | **0.04** | 71.41 | 24.07 |
| Watterson's $\theta$: $P$-value | 0.205 | 0.039 | **0.022\*** | 0.614 | 0.433 |
| Tajima's D: test statistic | 0.56 | -1.27 | -4.04 | 3.50 | 2.38 |
| Tajima's D: > 0 (%) | 51.93 | 39.59 | 27.39 | 73.21 | 69.72 |
| Tajima's D: $P$-value | 0.963 | 0.798 | 0.545 | 0.539 | 0.604 |
| $F_{ST}$: test statistic | 0.88 | 0.67 | 0.75 | -0.19 | -0.35 |
| $F_{ST}$: > 0 (%) | 77.94 | 88.78 | 87.07 | 39.64 | 26.83 |
| $F_{ST}$: $P$-value | 0.457 | 0.216 | 0.255 | 0.758 | 0.513 |
| 4-fold sites |  |  |  |  |  |
| $\pi$: test statistic | -0.13 | -0.04 | -0.16 | 0.01 | 0.06 |
| $\pi$: > 0 (%) | 11.68 | 16.31 | **0.55** | 59.01 | 86.68 |
| $\pi$: $P$-value. | 0.239 | 0.320 | **0.016\*** | 0.844 | 0.251 |
| Watterson's $\theta$: test statistic | -0.10 | -0.02 | -0.14 | 0.02 | 0.04 |
| Watterson's $\theta$: > 0 (%) | 10.54 | 23.08 | **0.16** | 68.92 | 84.08 |
| Watterson's $\theta$: $P$-value | 0.218 | 0.443 | **0.006\*\*** | 0.635 | 0.313 |
| Tajima's D: test statistic | -1.13 | -2.10 | 4.99 | -4.25 | 0.22 |
| Tajima's D: > 0 (%) | 46.02 | 31.94 | 78.49 | 21.43 | 51.46 |
| Tajima's D: $P$-value | 0.916 | 0.633 | 0.427 | 0.423 | 0.960 |
| $F_{ST}$: test statistic | -1.42 | -0.58 | -0.32 | -0.33 | -0.19 |
| $F_{ST}$: > 0 (%) | 9.24 | 6.90 | 31.19 | 31.71 | 37.14 |
| $F_{ST}$: $P$-value | 0.195 | 0.158 | 0.599 | 0.606 | 0.694 |

**Table 4.**
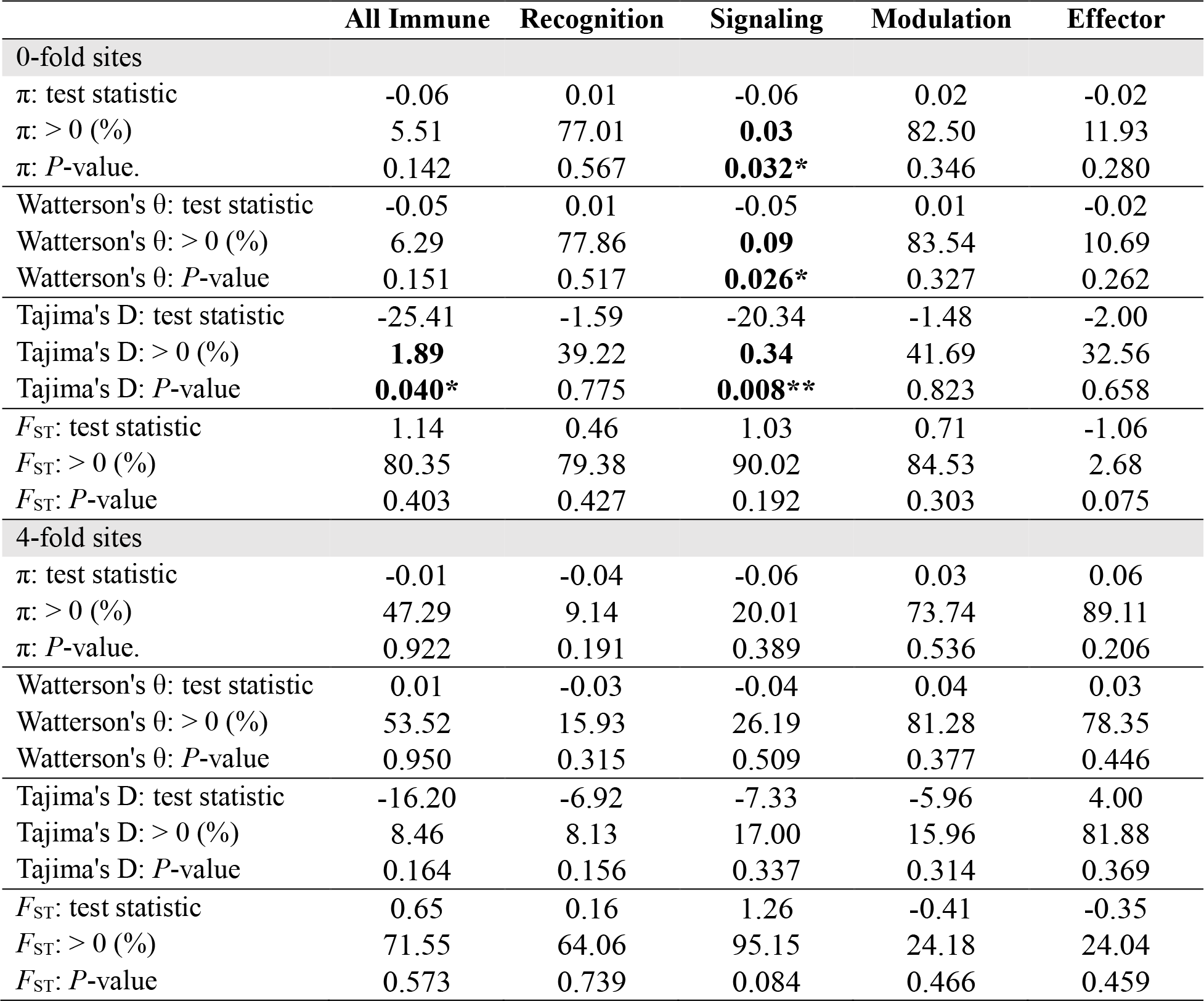
Population genetic statistics of immune genes in the Atlantic population using the paired-control approach. The upper half shows results based on the 0-fold sites and the lower half shows results based on the 4-fold sites. *F*_ST_ was compared to the North American population. “All immune” indicates the full immune gene set. In each statistic, the first row shows the test statistic of the immune gene group. The second row shows the proportion of 10,000 permutations in which the difference between the means of the immune gene group and the control set was positive. Percentages < 2.5% and > 97.5 % are labeled in bold. The third row shows the *P*-value. *P*-values < 0.05 are labeled in bold. Asterisks indicate: * < 0.05, ** < 0.01, *** < 0.001.

**Figure 6.**
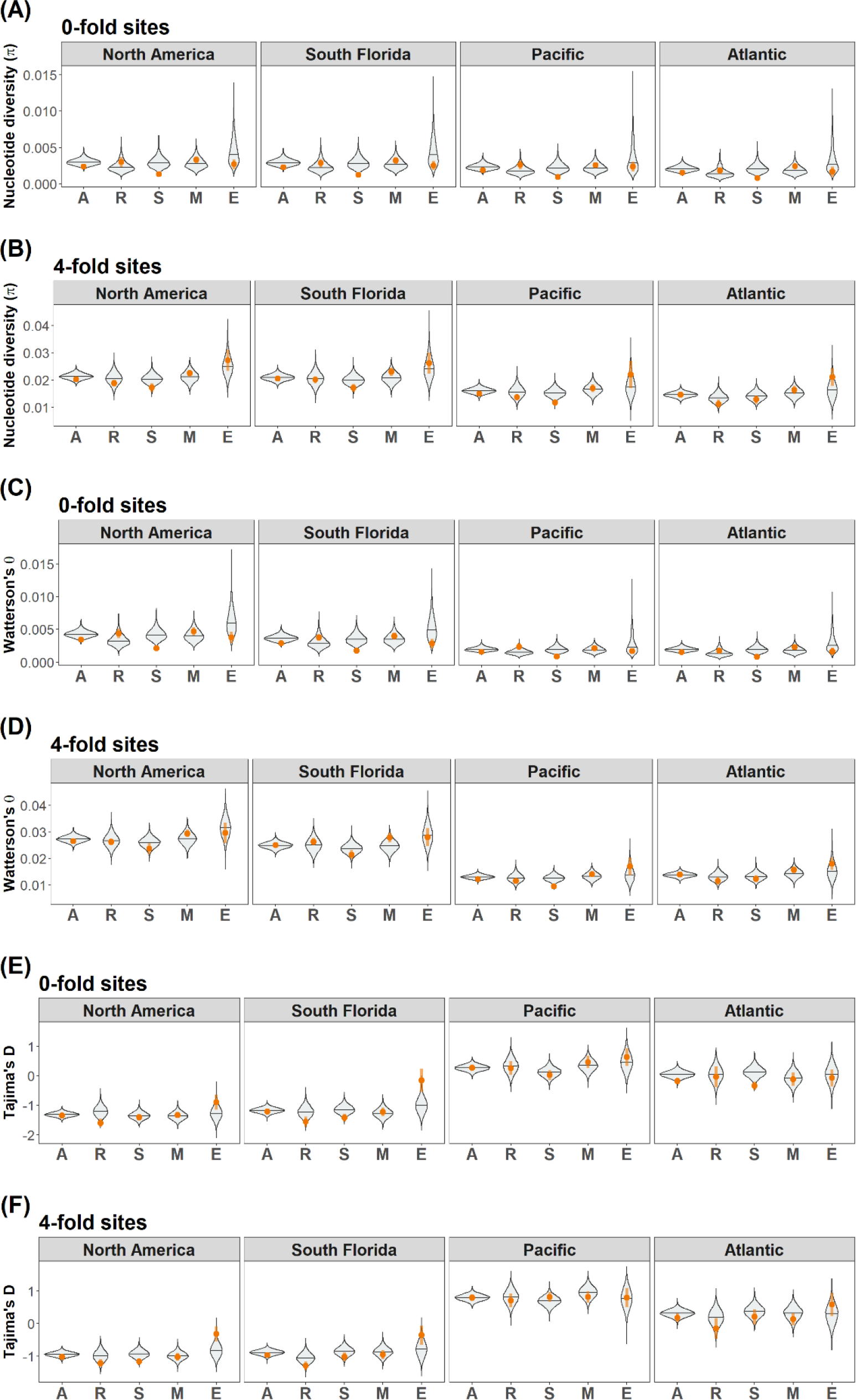
Population genetic statistics of immune genes in all four populations (North America, Florida, Pacific, and Atlantic) using the paired-control approach. 0-fold sites shown in (A), (C), and (E), and 4-fold sites shown in (B), (D), and (F). (A) and (B): Nucleotide diversity (**π**); (C) and (D): Watterson’s θ; (E) and (F): Tajima’s D. Each immune gene group was compared to selected pair-control sets. Violin plots show the distribution of the mean of each control set generated with 10, 000 permutations. The orange dots and vertical lines indicate mean ±1 SEM of the immune gene group of interest. X-axis represents immune gene groups: all immune genes (A), recognition genes (R), signaling genes (S), modulation genes (M), and effector genes (E).

As a group, immune genes were not under uniformly strong directional or balancing selection in any population, with one exception: in the Atlantic population, the 0-fold sites exhibited significantly lower Tajima’s D compared to the control genes, suggesting that, as a group, they experience increased directional selection (Tables 1–4 and Fig. 6). When considering genes of each functional class separately, there were differences in patterns not only between functional classes but also across populations. For recognition genes, the North America population showed a significantly lower Tajima’s D at the 0-fold sites than controls, but this was not found in any other population (Florida was marginally significant). For signaling genes, the Atlantic population showed a significantly lower Tajima’s D than controls at the 0-fold sites (Florida was marginally significant), but not at the 4-fold sites; in North America, Tajima’s D was significantly lower than controls at the 4-fold sites, but not at the 0-fold sites. For modulation genes, no significant differences to the controls were found across populations. For effector genes, both the North America and Florida populations displayed significantly higher Tajima’s D values compared to their controls: in North America, Tajima’s D was significantly higher than controls at the 4-fold sites, while in Florida the 0-fold sites showed higher Tajima’s D than controls.

Taken together, across all populations, immune genes as a group did not consistently exhibit significantly different levels of genetic variation and signatures of selection. Regarding genetic variation, a highly consistent pattern across populations was that the 0-fold sites of signaling genes showed significantly lower variation compared to control genes. There was also a trend for recognition and modulation genes to have greater variation than their respective controls. Regarding signatures of selection, the four populations exhibited moderately different patterns – there was no universal pattern across all populations. While effector genes displayed significantly higher Tajima’s D than controls, indicating balancing selection in some populations, recognition and signaling genes showed significantly lower Tajima’s D than their controls in some populations, indicating directional selection. Analyses based on all sites within each gene showed similar qualitative results (see supplemental information Tables S3-6 and Figs. S1-4).

#### B. Across-population analyses: population-level differentiations

We analyzed population differentiation using the ancestral population (*i.e*., North America) as the reference population (Tables 1–4 and Fig. 7). The full set of immune genes used in this study did not display any significant differentiation compared to control genes. Across each functional class, there were no universal differences. However, there was an overall non-significant trend across populations at 0-fold sites: recognition genes showed higher *F*_ST_ than controls while effectors displayed lower *F*_ST_ than controls. Between the Florida and the ancestral populations, recognition genes showed significantly greater *F*_ST_ than controls; between the Atlantic and the ancestral populations, effector genes showed marginally significantly lower *F*_ST_ than controls. Analyses based on all sites within each gene showed similar qualitative results (see supplemental information Tables S3-6 and Figs. S1-4).

**Figure 7.**
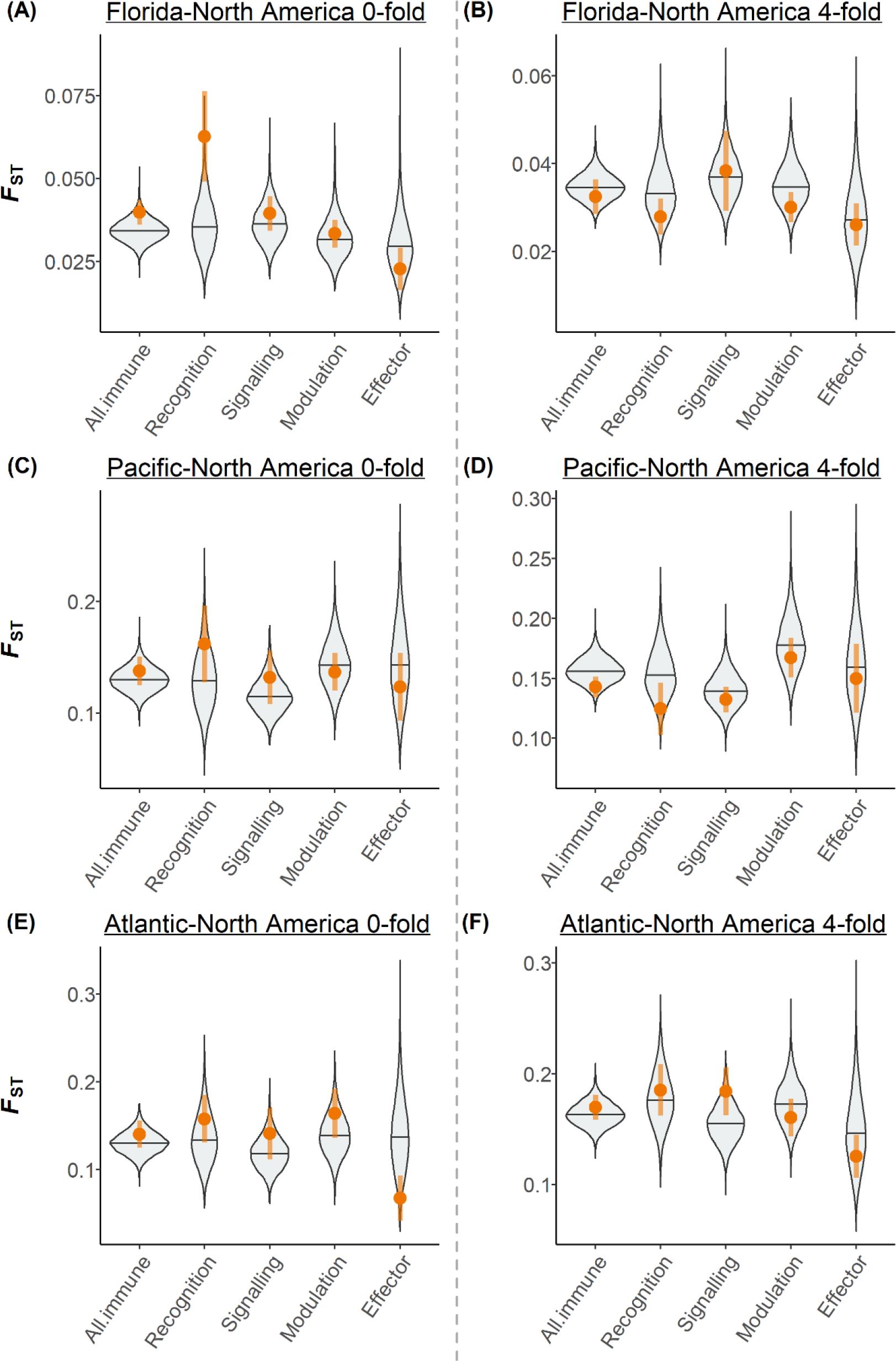
*F*_ST_ of immune genes in each derived population compared to the ancestral (North American) population using the paired-control approach. 0-fold sites shown in (A), (C), and (E), and 4-fold sites shown in (B), (D), and (F). (A)-(B): South Florida population (**π**); (C)-(D): Pacific population; (E)-(F): Atlantic population. Each immune gene group was compared to selected pair-control sets. Violin plots show the distribution of the mean of each control set generated with 10, 000 permutations. The orange dots and vertical lines indicate mean ±1 SEM of the immune gene group of interest.

### 3.3. Outlier analyses: access the patterns of outlier immune genes

We visualized Tajima’s D and *F*_ST_ results together to assess outlier patterns, considering both signatures of selection and differentiation among populations simultaneously. In Fig. 8, outliers that fall into different areas suggest different evolutionary scenarios. Different immune gene functional groups did not seem to show distinct differences in outlier patterns, but they differed greatly in the proportion of genes that were outliers, ranging from 14.3% to 31.6% at 0-fold sites and from 7.1% to 42.9% at 4-fold sites. We separated outlier genes into five categories based on their location in the 2D Tajima’s D-*F*_ST_ plot. We first compared whether the frequencies of outliers in each category (four outlier areas plus the central non-outlier area) differed across populations within each functional class. For the four functional classes, those frequencies did not differ significantly across populations at either the 0-fold or 4-fold sites (Table 5). Next, we compared whether the frequencies of outliers in each category differed across functional classes within each population. For the four populations, those frequencies did not differ significantly across functional classes at either the 0-fold or 4-fold sites (Table 6). In addition, we tested if immune genes, as one group, were disproportionally represented in genome-wide outliers, and found that they were not (see supplemental information Tables S7). Overall, our results indicate no statistically significant differences in outlier patterns across populations or functional classes.

**Table 5.** Contingency tables of Tajima’s D – *F*_ST_ outliers for each immune functional class. Numbers are counts of genes for each category. Q1 – Q4 categories represent the four quadrants shown in each Figure 8 plot (Q1 = top-right, Q2 = top-left, Q3 = bottom-left, Q4 = bottom-right). NS category represents non-outliers (i.e., the area within dotted gray lines). *P*-values from Fisher exact tests for each contingency table are shown in the last column. The North American population was not used because it was the reference group and did not have *F*_ST_ data.

| Gene class | Sites | Population | Q1 | Q2 | Q3 | Q4 | NS | $P$-value |
| --- | --- | --- | --- | --- | --- | --- | --- | --- |
| Recognition | 0-fold | Florida | 1 | 2 | 0 | 0 | 16 | 0.550 |
|  |  | Pacific | 2 | 0 | 0 | 0 | 17 |  |
|  |  | Atlantic | 0 | 1 | 0 | 0 | 18 |  |
| Signaling | 0-fold | Florida | 2 | 2 | 1 | 0 | 36 | 0.923 |
|  |  | Pacific | 1 | 1 | 2 | 0 | 37 |  |
|  |  | Atlantic | 0 | 2 | 2 | 0 | 37 |  |
| Modulation | 0-fold | Florida | 0 | 2 | 1 | 0 | 25 | 1.000 |
|  |  | Pacific | 1 | 1 | 0 | 0 | 26 |  |
|  |  | Atlantic | 1 | 1 | 1 | 0 | 25 |  |
| Effector | 0-fold | Florida | 0 | 0 | 0 | 2 | 12 | 0.761 |
|  |  | Pacific | 0 | 0 | 0 | 0 | 14 |  |
|  |  | Atlantic | 0 | 0 | 0 | 1 | 13 |  |
| Recognition | 4-fold | Florida | 0 | 0 | 0 | 0 | 19 | 0.766 |
|  |  | Pacific | 0 | 0 | 1 | 0 | 18 |  |
|  |  | Atlantic | 1 | 1 | 0 | 0 | 17 |  |
| Signaling | 4-fold | Florida | 0 | 2 | 4 | 1 | 34 | 0.182 |
|  |  | Pacific | 1 | 1 | 0 | 1 | 38 |  |
|  |  | Atlantic | 3 | 3 | 1 | 0 | 34 |  |
| Modulation | 4-fold | Florida | 0 | 0 | 0 | 0 | 28 | 1.000 |
|  |  | Pacific | 0 | 0 | 0 | 0 | 28 |  |
|  |  | Atlantic | 0 | 1 | 0 | 0 | 27 |  |
| Effector | 4-fold | Florida | 0 | 0 | 0 | 2 | 12 | 0.365 |
|  |  | Pacific | 0 | 0 | 0 | 0 | 14 |  |
|  |  | Atlantic | 0 | 1 | 1 | 1 | 11 |  |

**Table 6.** Contingency tables of Tajima’s D – *F*_ST_ outliers for each population. Numbers are counts of genes for each category. Q1 – Q4 categories represent the four quadrants shown in each Figure 8 plot (Q1 = top-right, Q2 = top-left, Q3 = bottom-left, Q4 = bottom-right). NS category represents non-outliers (i.e., the area within dotted gray lines). *P*-values from Fisher exact tests for each contingency table are shown in the last column. In the North American population, the analyses were based on only the Tajima’s D data. Q1 represents “right area” and Q2 represents “left area”. Q3 and Q4 were thus non-applicable.

| Population | Sites | Gene class | Q1 | Q2 | Q3 | Q4 | NS | $P$-value |
| --- | --- | --- | --- | --- | --- | --- | --- | --- |
| North America | 0-fold | Recognition | 0 | 0 | NA | NA | 19 | 1.000 |
|  |  | Signaling | 0 | 2 | NA | NA | 39 |  |
|  |  | Modulation | 0 | 1 | NA | NA | 27 |  |
|  |  | Effector | 0 | 0 | NA | NA | 14 |  |
| Florida | 0-fold | Recognition | 1 | 2 | 0 | 0 | 16 | 0.374 |
|  |  | Signaling | 2 | 2 | 1 | 0 | 36 |  |
|  |  | Modulation | 0 | 2 | 1 | 0 | 25 |  |
|  |  | Effector | 0 | 0 | 0 | 2 | 12 |  |
| Pacific | 0-fold | Recognition | 2 | 0 | 0 | 0 | 17 | 0.820 |
|  |  | Signaling | 1 | 1 | 2 | 0 | 37 |  |
|  |  | Modulation | 1 | 1 | 0 | 0 | 26 |  |
|  |  | Effector | 0 | 0 | 0 | 0 | 14 |  |
| Atlantic | 0-fold | Recognition | 0 | 1 | 0 | 0 | 18 | 0.819 |
|  |  | Signaling | 0 | 2 | 2 | 0 | 37 |  |
|  |  | Modulation | 1 | 1 | 1 | 0 | 25 |  |
|  |  | Effector | 0 | 0 | 0 | 1 | 13 |  |
| North America | 4-fold | Recognition | 0 | 0 | NA | NA | 19 | 0.105 |
|  |  | Signaling | 0 | 2 | NA | NA | 39 |  |
|  |  | Modulation | 0 | 1 | NA | NA | 27 |  |
|  |  | Effector | 2 | 0 | NA | NA | 12 |  |
| Florida | 4-fold | Recognition | 0 | 0 | 0 | 0 | 19 | 0.092 |
|  |  | Signaling | 0 | 2 | 4 | 1 | 34 |  |
|  |  | Modulation | 0 | 0 | 0 | 0 | 28 |  |
|  |  | Effector | 0 | 0 | 0 | 2 | 12 |  |
| Pacific | 4-fold | Recognition | 0 | 0 | 1 | 0 | 18 | 0.833 |
|  |  | Signaling | 1 | 1 | 0 | 1 | 38 |  |
|  |  | Modulation | 0 | 0 | 0 | 0 | 28 |  |
|  |  | Effector | 0 | 0 | 0 | 0 | 14 |  |
| Atlantic | 4-fold | Recognition | 1 | 1 | 0 | 0 | 17 | 0.489 |
|  |  | Signaling | 3 | 3 | 1 | 0 | 34 |  |
|  |  | Modulation | 0 | 1 | 0 | 0 | 27 |  |
|  |  | Effector | 0 | 1 | 1 | 1 | 11 |  |

**Figure 8.**
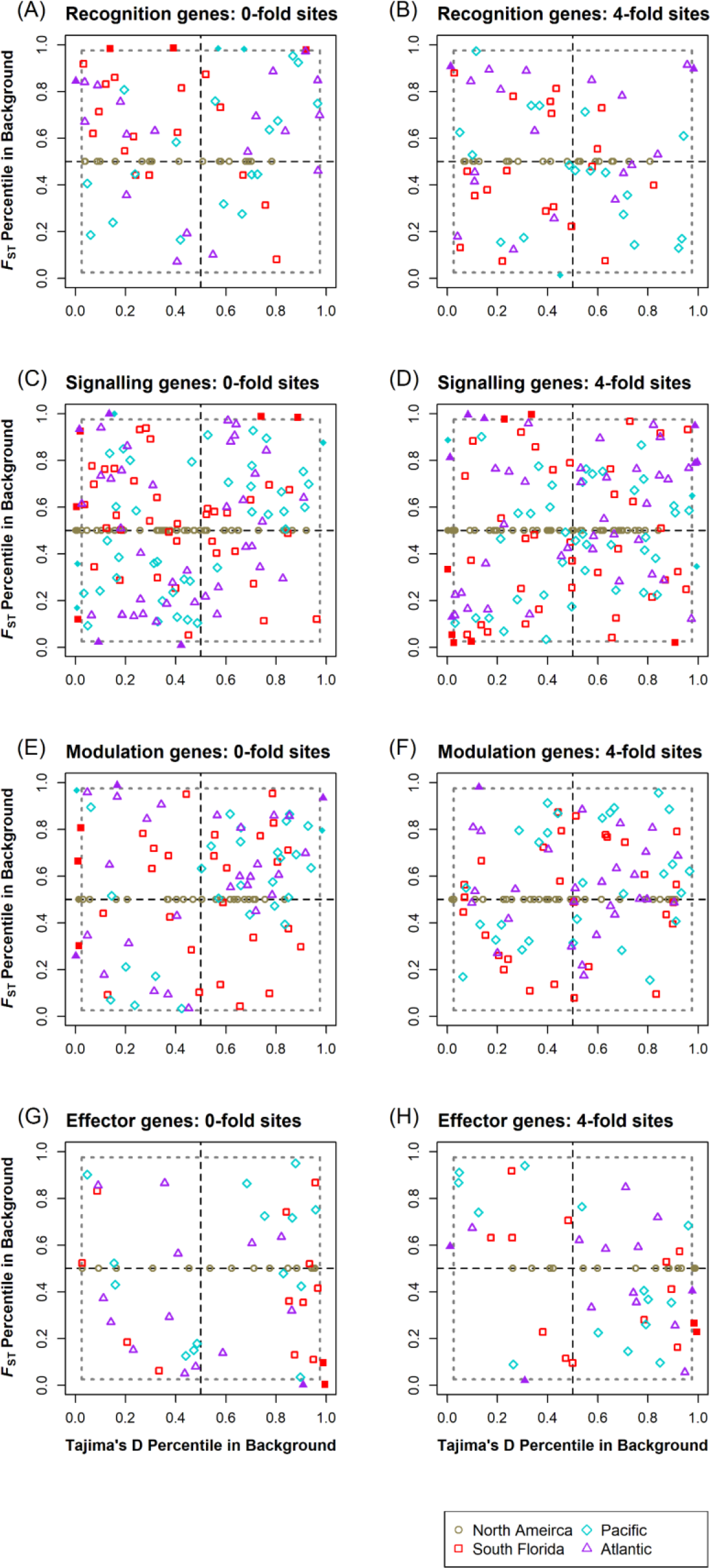
Tajima’s D – *F*_ST_ plots of the four immune gene functional classes. 0-fold sites shown in (A) – (D) and 4-fold sites shown in (E) – (H). (A) and (E): recognition (N = 19; 57.9% outlier in 0-fold; 31.6% outlier in 4-fold); (B) and (F): signaling (N = 41; 36.6% outlier in 0-fold; 53.7% outlier in 4-fold); (C) and (G): modulation (N = 28; 46.6% outlier in 0-fold; 17.9% outlier in 4-fold); (D) and (H): effector (N = 14; 64.3% outlier in 0-fold; 57.1% outlier in 4-fold). In each plot, populations were labeled in different colors and shapes. One dot represents one immune gene in one population, shown as their percentile in the genome background. Solid dots are outliers. Outliers were defined as < 2.5^th^ percentile or > 97.5^th^ percentile of the genome background. Dotted black lines indicate the median of genome background in each of the two measures, dividing the plot into four quadrants. Dotted gray lines indicate the boundaries of outlier areas. All data from the North American population, the reference population for *F*_ST_, were plotted on the y = 0.5 horizontal line since they do not have *F*_ST_ results.

We identified individual immune genes that were genome outliers based on 0-fold sites and summarized their statistics across populations (Table 7–8). Some genes exhibited distinct patterns across populations, as indicated by being outliers at different ends of the statistics. For example, *Pellino*, which belongs to the Toll pathway, was under directional selection (low Tajima’s D) in the Florida population, while under balancing selection (high Tajima’s D) in the Pacific population. One CLIP serine protease was under directional selection (low Tajima’s D) in the Pacific population, while under balancing selection (high Tajima’s D) in the Atlantic population. In addition, some of the patterns observed for *F*_ST_ outliers were population-specific – only shown in one population but not the others. Two out of three *Nimrod* genes were identified as *F*_ST_ outliers in the Pacific population compared to the ancestral population, and all of them showed higher differentiation (high *F*_ST_). Two out of seven Scavenger receptor (SCR) genes were identified as *F*_ST_ outliers in the Florida population, and all of them showed higher differentiation (high *F*_ST_). In contrast, some genes were identified as outliers in half of the populations in the same direction. For instance, one Toll-like receptor and *DOMELESS* were under directional selection (low Tajima’s D) in both the North America and Florida populations at the 0-fold sites; one *Attacin*-like gene showed lower differentiation (low *F*_ST_) in the Florida and Atlantic populations. However, no immune genes were consistently identified as outliers across all populations based on either Tajima’s D or *F*_ST_. Three genes were identified as outliers based on both Tajima’s D and *F*_ST_: Myeloid differentiation primary response 88 (*MyD88*), Protein inhibitor of activated STAT (*PIAS*), and one Attacin-like gene. *PIAS* showed a general trend of lower Tajima’s D and lower *F*_ST_, suggesting that it might be evolutionarily constrained. *MyD88* showed a general trend of higher *F*_ST_ and was an outlier in the Florida population. Also, in the Atlantic population, *MyD88* was a Tajima’s D outlier, indicating directional selection. The Attacin-like gene showed a general trend of higher Tajima’s D and was an outlier in the Florida population, indicating balancing selection. Also, it was an *F*_ST_ outlier in the Atlantic and Florida populations, indicating low differentiation. In summary, although our results did not reveal clear patterns of outliers based on functional groups or populations, individual outlier genes were identified. These results suggest that immune genes undergo individual evolutionary trajectories, and these trajectories vary across populations.

**Table 7.** Summary of immune genes that are outliers according to Tajima’s D at the 0-fold sites. Outliers were defined as < 2.5^th^ percentile or > 97.5^th^ percentile of the genome background. A gene was reported as an outlier when it met the criteria in at least one of the populations. The Tajima’s D value of the 0-fold sites and the 4-fold sites of each outlier gene are shown. Values that are less than or equal to the genome median (i.e., 50^th^ percentile) are underscored; values that are greater than the genome median are in italics. Values that are outliers are in bold. Genes that are reported as outliers in both Tajima’s D and *F*_ST_ (Table 8) are colored in red. A Tajima’s D value close to 0 indicates neutrality. A more negative Tajima’s D value represents an excess of low-frequency polymorphisms than expectation, which indicates directional selection or population expansion. A more positive Tajima’s D value represents low levels of both low- and high-frequency polymorphisms, which indicates balancing selection or population contraction.

| Gene name | Gene number | Functional class | <u>North America</u> |  | <u>Florida</u> |  | <u>Pacific</u> |  | <u>Atlantic</u> |  |
| --- | --- | --- | --- | --- | --- | --- | --- | --- | --- | --- |
|  |  |  | 0-fold | 4-fold | 0-fold | 4-fold | 0-fold | 4-fold | 0-fold | 4-fold |
| BGRP-like | DPOGS212941 | Recognition | <u>-1.74</u> | <u>-1.93</u> | <u>-1.90</u> | <u>-2.13</u> | *0.59* | <u>0.05</u> | **<u>-2.66</u>** | <u>-0.72</u> |
| CLIP-like | DPOGS204835 | Modulation | <u>-2.42</u> | **<u>-2.28</u>** | <u>-1.79</u> | <u>-1.72</u> | *0.07* | <u>0.40</u> | **<u>-2.72</u>** | *0.06* |
| CLIP-like | DPOGS204146 | Modulation | *-1.13* | <u>-1.17</u> | *-0.37* | <u>-1.30</u> | **<u>-2.08</u>** | <u>-0.14</u> | **<u>2.67</u>** | <u>-0.20</u> |
| CLIP-like | DPOGS208169 | Modulation | **<u>-2.54</u>** | *-0.91* | <u>-2.21</u> | *-0.67* | <u>-0.67</u> | <u>-0.19</u> | <u>-0.26</u> | *1.90* |
| CLIP-like | DPOGS211355 | Modulation | <u>-2.01</u> | <u>-1.81</u> | **<u>-2.67</u>** | <u>-1.56</u> | <u>-0.53</u> | <u>-0.03</u> | <u>-0.32</u> | *0.74* |
| CLIP-like | DPOGS214570 | Modulation | <u>-1.43</u> | *-0.79* | *-0.21* | *0.26* | **<u>2.44</u>** | *0.72* | <u>-0.69</u> | <u>-0.78</u> |
| CLIP-like | DPOGS206224 | Modulation | <u>-2.30</u> | <u>-1.31</u> | **<u>-2.64</u>** | <u>-1.78</u> | *-0.06* | <u>-0.39</u> | <u>-0.98</u> | <u>-1.49</u> |
| CLIP-like | DPOGS205206 | Modulation | *-0.81* | <u>-1.28</u> | **<u>-2.59</u>** | <u>-1.62</u> | <u>-0.92</u> | *1.17* | <u>-1.25</u> | *0.04* |
| Toll-like receptors | DPOGS205295 | Signaling - Toll | **<u>-2.60</u>** | *-0.80* | **<u>-2.77</u>** | *0.31* | <u>-0.97</u> | *0.88* | *0.33* | *1.19* |
| **MyD88** | **DPOGS205936** | **Signaling - Toll** | *-0.29* | *-0.61* | *-0.07* | *-0.43* | *0.99* | **<u>2.50</u>** | **<u>-2.21</u>** | **<u>-2.45</u>** |
| Pellino | DPOGS214647 | Signaling - Toll | *-0.95* | <u>-1.88</u> | **<u>-2.60</u>** | <u>-1.37</u> | **<u>2.54</u>** | *0.83* | <u>-0.46</u> | <u>-1.09</u> |
| Hem | DPOGS208954 | Signaling - JNK | <u>-1.42</u> | *-0.80* | <u>-1.54</u> | <u>-1.34</u> | **<u>-1.98</u>** | <u>0.70</u> | <u>-0.62</u> | <u>-0.06</u> |
| **PIAS** | **DPOGS214325** | **Signaling - JAK-STAT** | <u>-1.58</u> | <u>-1.09</u> | <u>-1.40</u> | <u>-1.37</u> | **<u>-1.93</u>** | *0.87* | <u>-0.30</u> | *1.57* |
| DOMELESS | DPOGS200349 | Signaling - JAK-STAT | **<u>-2.79</u>** | *-0.18* | **<u>-2.68</u>** | *-0.08* | <u>-1.24</u> | *1.70* | <u>-0.83</u> | *0.73* |
|  |  |  | 0-fold | 4-fold | 0-fold | 4-fold | 0-fold | 4-fold | 0-fold | 4-fold |
| Attacin-Like | DPOGS213997 | Effector | *0.23* | *-0.22* | ***2.03*** | <u>*-1.00*</u> | *2.06* | *0.82* | *1.83* | *0.52* |
| PPO-like | DPOGS200017 | Effector | *-0.74* | <u>*-1.30*</u> | ***1.62*** | <u>*-1.19*</u> | <u>*-1.48*</u> | <u>*-0.42*</u> | <u>*-0.38*</u> | <u>***-2.43***</u> |

**Table 8.** Summary of immune genes that are outliers according to *F*_ST_ at the 0-fold sites. Outliers were defined as < 2.5th percentile or > 97.5^th^ percentile of the genome background. A gene was reported as an outlier when it met the criteria in at least one of the population pairs. The *F*_ST_ value of the 0-fold sites and the 4-fold sites of each outlier gene are shown. Values that are less than or equal to the genome median (i.e., 50^th^ percentile) are underscored; values that are greater than the genome median are in italics. Values that are outliers are in bold. Genes that are reported as outliers in both Tajima’s D (Table 7) and *F*_ST_ are colored in red. *F*_ST_ is a measure of population differentiation due to genetic structure, with a value ranging from 0 to 1. An *F*_ST_ value equals to zero indicates no differentiation. An *F*_ST_ value equals to one indicates complete differentiation; different alleles are fixed in different populations.

| Gene name | Gene number | Functional class | <u>Florida – North America</u> |  | <u>Pacific – North America</u> |  | <u>Atlantic – North America</u> |  |
| --- | --- | --- | --- | --- | --- | --- | --- | --- |
|  |  |  | 0-fold | 4-fold | 0-fold | 4-fold | 0-fold | 4-fold |
| BGRP-like | DPOGS212940 | Recognition | ***0.14*** | <u>0.02</u> | *0.20* | <u>0.12</u> | *0.14* | <u>0.03</u> |
| Class B-like SCR | DPOGS203180 | Recognition | ***0.12*** | <u>0.01</u> | *0.17* | <u>0.10</u> | *0.15* | <u>0.11</u> |
| Other SCR | DPOGS214397 | Recognition | ***0.15*** | *0.05* | <u>0.07</u> | *0.20* | *0.46* | *0.30* |
| NIM-like | DPOGS210210 | Recognition | *0.06* | *0.04* | ***0.48*** | *0.20* | *0.28* | <u>0.09</u> |
| NIM-like | DPOGS210211 | Recognition | *0.04* | *0.05* | ***0.50*** | *0.43* | *0.25* | *0.30* |
| CLIP-like | DPOGS215098 | Modulation | *0.04* | *0.03* | *0.15* | *0.15* | ***0.57*** | ***0.48*** |
| SPZ-like | DPOGS209810 | Signaling - Toll | <u>0.02</u> | **<u>-0.01</u>** | <u>0.03</u> | <u>0.03</u> | **<u>0.00</u>** | <u>0.06</u> |
| MyD88 | DPOGS205936 | Signaling - Toll | ***0.14*** | *0.10* | *0.27* | *0.17* | *0.35* | *0.24* |
| JNK | DPOGS213169 | Signaling - JNK | ***0.16*** | ***0.26*** | <u>0.06</u> | *0.15* | <u>0.02</u> | <u>0.03</u> |
| PIAS | DPOGS214325 | Signaling - JAK-STAT | <u>0.00</u> | <u>0.01</u> | <u>0.05</u> | *0.21* | **<u>-0.01</u>** | <u>0.08</u> |
| Stat | DPOGS212956 | Signaling - JAK-STAT | *0.04* | *0.06* | ***0.86*** | *0.29* | ***0.80*** | ***0.64*** |
| Attacin-Like | DPOGS213997 | Effector | **<u>-0.02</u>** | <u>0.00</u> | *0.17* | *0.21* | **<u>-0.02</u>** | *0.15* |

The analysis of outliers supports our notion that the complex evolutionary pressures have resulted in different patterns of selection on individual genes in the different populations, involving a wide variety of biological processes and targets. A few genes that showed high population differentiation (*F*_ST_ outliers at the upper end) are involved in cellular immune processes, such as phagocytosis. Two SCR genes showed high differentiation only in the Florida population, while two *nimrod* genes showed high differentiation only in the Pacific population. Notably, the *Nimrod* gene family is involved in recognizing foreign object for phagocytosis, which likely has direct interactions with pathogens (Estévez-Lao & Hillyer, 2014; Kurucz et al., 2007; Somogyi, Sipos, Pénzes, & Andó, 2010). Several of the outlier genes either belong to or interact with the Toll signaling pathway. For instance, two outlier genes encode Beta-1,3-glucan recognition proteins (BGRPs), both of which recognize bacterial and/or fungal signals and are known to activate the toll signaling cascade in *Drosophila* (Kim et al., 2000). One of them is involved in activation of the phenoloxidase cascade (Matskevich, Quintin, & Ferrandon, 2010), while the other one leads to signal transmission that induces the expression of AMPs such as cecropin and attacin (Kim et al., 2000). Some members of the Toll pathway, such as *spaetzle*, *Pellino*, and *MyD88*, were identified as outliers. *MyD88*, which is involved in regulating AMPs in *Drosophila* (Tauszig-Delamasure, Bilak, Capovilla, Hoffmann, & Imler, 2002). Attacins, which are AMPs against Gram-negative bacteria, are regulated mostly by the IMD pathway but also known to have some interactions with the Toll pathway (Tanji, Hu, Weber, & Ip, 2007).

## 4 DISCUSSION

Our results demonstrate that immune genes, as one group, do not exhibit uniform patterns of selection, differentiation, or high genetic variation; different function classes show different patterns. Monarchs recently spread around the world via three main dispersal events (Pierce et al., 2014; Zhan et al., 2014). During these colonization processes, they have encountered different ecological conditions that are likely to drive the evolution of immune genes. Our results show that patterns of evolutionary change in immune genes of different functional groups vary to some extent across populations, suggesting that populations might not be under a uniform selection regime. This is further supported by assessing individual genes that are genome outliers, as some of them exhibit distinct differences across populations.

### 4.1 Population genomic patterns and adaptive evolution across different functional classes

A limited body of work has demonstrated that different components of the canonical insect immune system can face distinct selection pressures. Genes encoding proteins in the core signaling pathways, for example, have been shown to be more functionally constrained (Sackton et al., 2007). Similarly, low genetic variation in signaling genes is one of the most consistent patterns found in monarchs – signaling genes showed significantly lower genetic variation than control genes in all the populations studied. Most likely, this reflects the increased removal of deleterious alleles among these loci. The DFEs of signaling genes also points to this phenomenon, indicating a much larger proportion of strongly deleterious variants among new mutations relative to control genes. Broadly increased purifying selection can also help to explain the greater α value, which indicates a higher proportion of adaptive amino acid substitutions between species. If most new mutations are removed by purifying selection, then any divergence observed should primarily reflect adaptation, not neutral divergence (*i.e*., drift), even though the absolute amount of divergence might be relatively low. Indeed, such increased purifying selection, if consistent over long periods of time, should reduce overall divergence between species. However, while signaling genes do have reduced average divergence relative to controls, this difference is not significant. Thus, it is possible that the strong purifying selection we observe in *D. plexippus* is a relatively recent phenomenon that manifests patterns in population diversity but not yet at the level of species divergence. Further population genetic analysis in other *Danaus* species would be required to assess whether there are long-term patterns of selection for this group of butterflies. Given the broad finding of functional constraint in other distantly related species, such variability in evolutionary pressures among signaling genes between closely related species is an intriguing possibility.

In striking contrast to signaling genes, modulation genes show a consistent pattern of increased diversity. While nucleotide diversity is only somewhat elevated, and not significantly so, interspecific divergence is greatly increased. One good explanation for these patterns is that modulation genes experience relaxed selection compared to controls. This idea fits well with patterns in the DFEs, which indicates notably more neutral variants and fewer strongly deleterious variants among new mutations among modulation genes. The relatively rapid divergence of modulation genes due to fixation of neutral or weakly deleterious mutations can explain the reduced α value. Taken together, signaling and modulation genes both exhibited consistent evolutionary patterns across populations, suggesting that the selection regime on these two functional classes might not differ strongly across populations.

Signaling genes and modulation genes are sometimes considered as one functional class (*e.g.*, Waterhouse et al., 2007), but our results show distinct differences in genetic diversity. Signaling genes, especially those within the Toll, IMD, JAK-STAT, and JNK pathways, are well-characterized for their function. However, relatively little is known about the functional roles of modulation genes, most of which are CLIP serine proteases (Lemaitre & Hoffmann, 2007). Our results suggest that signaling and modulation genes likely have different functional roles, as they exhibit notably distinct patterns of selection.

In contrast, genes that encode proteins that have direct interactions with pathogens, such as recognition and effector genes, have been shown to evolve more rapidly as they are more likely targets of host-pathogen coevolutionary arms races (Sackton et al., 2007). For effector genes, recent studies have demonstrated signatures of balancing selection in some taxa, especially for AMPs (Chapman et al., 2018; Unckless et al., 2016; Unckless & Lazzaro, 2016). Similarly, in monarchs, effector genes show notable evidence of balancing selection. Specifically, Tajima’s D is elevated, at least in some populations. However, this elevated Tajima’s D occurs without a clear signal of broadly elevated heterozygosity, which would be expected in many scenarios involving balancing selection. This discrepancy in observed patterns might result if a few effector genes show strongly balanced patterns, contributing substantially to greater Tajima’s D but less so to average variation across effector loci. Anticipating or interpreting the DFE under balancing selection is not straightforward (Connallon & Clark, 2015). Yet it is very clear that the DFE is qualitatively distinct between effectors and their controls, as well as the other classes of immune genes: there appear to be many fewer new mutations showing strongly deleterious effects. Subsequently, a greater proportion of these less-deleterious variants reach the intermediate frequencies associated with balancing selection.

We observed considerable differences in patterns of selection across populations on effector genes. Specifically, signatures of balancing selection were observed in the North America and Florida populations but not in the Pacific and Atlantic populations. One possible interpretation is that this pattern reflects a shift in selective regime among populations. When monarchs dispersed to distant locations across the Pacific and Atlantic oceans, the selection regimes shifted toward either directional selection or were relaxed, leading to a loss of selective signal in these two populations. Alternatively, the selective regime may be constant, but demographic effects, including bottlenecks and other non-equilibrium effects, are masking the signal. Specifically, bottleneck effects, which the Pacific and Atlantic populations have experienced (Pierce et al., 2014; Zhan et al., 2014), can skew allele frequencies. The effect of skewed allele frequencies due to bottlenecks can obscure the signal of balancing selection. In a more extreme scenario, one of the selected variants could be entirely removed by bottlenecks so that the balanced polymorphism cannot be restored after the population recovered. Even though we tried to account for demographic effects by using a paired-control approach, there is still a possibility that we have reduced resolution in the derived populations due to demographic effects.

Evolutionary analyses of immune genes in other species, particularly *Drosophila*, indicate that recognition genes have the strongest evidence for adaptive evolution among immune functional groupings (McTaggart et al., 2012; Sackton et al., 2007). By comparison, there was distinctly mixed evidence for strong selection among recognition genes in monarchs. In North America (and Florida), Tajima’s D was notably reduced relative to controls for both 0-fold and 4-fold sites, though without much reduction in 4-fold heterozygosity, and even a modest increase for 0-fold heterozygosity. If this pattern reflects recent selective sweeps among some recognition genes for these populations, it is likely a narrow range of parameters that would produce such skewed distributions of diversity (*i.e.*, Tajima’s D) without also affecting the amount of diversity (i.e. heterozygosity). Nonetheless, recurring adaptation among recognition genes could also explain the significantly elevated Dn/Ds observed in divergence to *D. gilippus*. Alternatively, this could result from relaxed constraint, as we argued above for modulators. Also, like modulators, the DFE of recognition genes suggests relatively fewer strongly selected variants compared to controls, and α is also lower. The mixed signals for selection in recognition loci also play out among patterns of population differentiation. The 0-fold *F*_ST_ between North American and Florida populations is strongly elevated relative to controls; a similar but less extreme signal occurs for Pacific vs. North America. While this could be interpreted as evidence for local adaptation among these distinct populations, no such pattern was observed among linked 4-fold sites, which might be expected to show the same pattern due to background selection. These contrasting patterns among the different analytical components employed here are not easily synthesized into a single coherent biological interpretation for recognition loci; a more detailed, gene-by-gene analysis may be required to resolve many of these discrepancies.

We also observed differences in patterns of selection across populations on recognition genes. Specifically, significant signatures of directional selection were observed in the North America population, but they were only marginally significant in the Florida population, and not significant in the Pacific and Atlantic populations. Intriguingly, this pattern across populations is similar to what was observed for the balancing selection on effector genes. One possible interpretation is that this pattern reflects a shift in selective regime among populations. That is, the differences reflect local adaption to pathogens. Alternatively, the selective regime may be constant, but demographic effects are masking the signal. Bottleneck effects can exert similar effects as selective sweeps, removing rare alleles, but acting across the entire genome instead. The removal of rare alleles can result in a disproportional loss of genetic variation on loci with high-and intermediate-level polymorphisms compared to loci under directional selection, which already have lower polymorphism. That is, bottlenecks can result in a disproportional loss of genome-wide genetic variation compared to loci under directional selection. Similar to directional selection, selection sweeps can result in a low Tajima’s D value by removing rare alleles (Nielsen & Slatkin, 2013). Therefore, although we tried to account for demographic effects in our analyses, there is still a possibility that we have a reduced resolution in the derived populations.

Overall, our results and those of previous studies on *Drosophila melanogaster* and *Pieris napi* (Early et al., 2017; Keehnen et al., 2018) highlight that it may be common for different components of the insect canonical immune system to have different evolutionary trajectories. A common trend among the three taxa is that genes within signaling pathways show lower levels of genetic variation, genes involve in recognition show higher levels of population differentiation in some scenarios (Early et al., 2017; Keehnen et al., 2018), and that genes encoding effector molecules (especially AMPs) show signatures of balancing selection (Chapman et al., 2018; Keehnen et al., 2018; Unckless et al., 2016). The emergence of these common patterns across insect species that differ considerably in life histories and taxonomy suggests that there may be some general evolutionary patterns among insect immune genes.

### 4.2 Ecological differences among populations and their potential consequences for immune gene evolution

Ecological factors that vary across populations affect the strength and type of selection and can therefore lead to local adaptation (Eizaguirre et al., 2012). When monarchs dispersed around the world, they experienced novel ecological conditions, likely resulting in differential selection across populations. First, different populations face different pathogen pressures. In monarchs, the most common and best-understood parasite is the virulent specialist protozoan parasite *Ophryocystis elektroscirrha*, which occurs at low prevalence in the ancestral North American population but at much greater prevalence in tropical and sub-tropical locations that monarchs colonized during their worldwide dispersal (Altizer & de Roode, 2015), resulting in greater parasitism risk and possibly stronger selection of monarch immunity. Second, although North American monarchs migrate thousands of kilometers to overwinter in Central Mexico, the derived populations that established during world-wide dispersal have become non-migratory (Zhan et al., 2014). This loss of migration is likely partly responsible for the increased parasite prevalence in derived populations. In North America, the strenuous annual migration weeds out heavily infected monarchs, thus reducing parasite prevalence. In non-migratory populations, this seasonal break on parasite transmission has been eliminated, leading to greater transmission and prevalence (Altizer & de Roode, 2015; Altizer, Hobson, Davis, De Roode, & Wassenaar, 2015; Bartel, Oberhauser, de Roode, & Altizer, 2011). Although this greater prevalence may select for greater immunity, it is also possible that the lack of a migratory phase, and the accompanying lack of a generation that needs to survive for long periods of time as it flies thousands of kilometers, results in less investment in immunity. Third, while the majority of North American monarchs utilize *Asclepias syriaca* (common milkweed) as their larval host plant, monarchs in newly colonized populations rely on other species, including *A. curassavica*, *A. fruticosa*, and *A. physocarpa*. Notably, these species have greater concentrations of cardenolides (secondary toxic compounds), which have been shown to reduce *O. elektroscirrha* infection, growth and virulence (Gowler, Leon, Hunter, & de Roode, 2015; Sternberg et al., 2012; Tao, Hoang, Hunter, & de Roode, 2016). The use of such medicinal compounds could in theory relax selection on immune genes, especially when immune responses are costly (de Roode, Lefèvre, & Hunter, 2013; Evans et al., 2006; Gerardo et al., 2010; Parker et al., 2011). Finally, while we know most about parasitism by *O. elektroscirrha*, monarchs are undoubtedly challenged by a suite of pathogens that vary in presence and prevalence across populations. These differences in disease pressure undoubtedly shape the evolution of monarch immune defenses.

The different ecological conditions experienced by monarchs as they dispersed around the world do not act in isolation, resulting in a complex mosaic of factors that simultaneously select for greater or lesser investment in immunity. Furthermore, the evolutionary patterns of immune gene evolution also may be influenced by demographic history and stochastic processes. In our analyses, immune genes as a group did not display consistent patterns across populations. For instance, directional selection on recognition genes, which indicates an excess of rare alleles, was only seen in the North America population (Florida was marginally significant). Furthermore, different immune genes were outliers in different populations. This difference among populations could in part be driven by genetic drift rather than differential selection; however, few immune genes were identified as outliers in multiple populations with strikingly different patterns. For example, *Pellino*, which belongs to the Toll pathway, showed an excess of rare alleles in the Florida population (directional selection) but showed maintenance of multiple alleles at moderate frequency (balancing selection) in the Pacific population, indicating that the selection forces between these two populations are very different.

## 5 CONCLUSIONS

In summary, our results demonstrate that immune genes as a whole are not under uniform patterns of selection or differentiation compare to the genome background. Different components of the immune system exhibit different evolutionary patterns. Signaling genes exhibit consistently low levels of genetic variation across populations and between the two *Danaus* species, indicating they are likely very constrained, while modulation genes exhibit the opposite pattern - signatures of relaxed selection. In contrast, effector and recognition genes exhibit less consistent patterns across populations. In some populations, effector genes exhibit signatures of balancing selection, while recognition genes exhibit directional selection and population differentiation. We find some clear differences among populations for individual genes that are genomic outliers, suggesting that immune genes undergo individual evolutionary trajectories. To a lesser extent, we also find some population-specific differences when considering each functional class separately. These results support the hypothesis that monarch populations do not face uniform selection pressures on immune genes.

The identification of immune genes that are under differential selection in monarch populations opens the way for further functional and ecological characterization. In particular, population-specific patterns indicate a possibility of local adaptation, and functional characterization is needed to understand the phenotypic effects of different alleles of immune genes, especially as they relate to important ecological factors, such as the prevalence of *O. elektroscirrha* and the use of medicinal milkweeds. Such functional characterization is also needed because several insect immune genes, especially signaling genes, have pleiotropic functions in immunological and non-immunological processes (Lemaitre & Hoffmann, 2007). Therefore, evolutionary patterns on those genes may not be solely driven by selection pressures on immunity.

## ACKNOWLEDGMENTS

We thank William J. Palmer for providing immune gene annotation resources for *Danaus plexippus*. We thank Robert L. Unckless, Joanna Chapman, Erika Diaz-Almeyda, Amanda A. Peirce, Yaw Kumi-Ansu, Venkat Talla, and the KU EEB-genetics group for helpful discussion on this research, and Robert A. Pettit III and Yu-Hui Lin for helpful guidance and discussion on bioinformatics computing. Analyses were performed on resources provided by the University of Kansas Information and Telecommunication Technology Center. This work was supported by National Science Foundation (NSF) grant IOS-1557724 to JCdR and NMG, and NSF grant DEB-1457758 to JRW.

## DATA ACCESSIBILITY

All sequence data were previously publicly available (Zhan et al., 2014). Custom analysis scripts can be found in the following GitHub repository: https://github.com/WaltersLab/Monarch_immune_evolution

## AUTHOR CONTRIBUTIONS

WHT, JCdR, and NMG conceived and designed the study, with JRW providing additional input. WHT performed the population genetic analyses and outlier analyses, and AJM performed the between-species analyses. JCdR, NMG and JRW supervised the bioinformatic and population genetic analyses. WHT wrote the initial draft of the manuscript and all authors have edited the manuscript.

