## Supplemental Information for "Population genomics reveals complex patterns of immune gene evolution in monarch butterflies (*Danaus plexippus*)"

### Table of Contents:


| | |
|---|---|
| **Table S1.** *Danaus* samples used in this study | Page 2 |
| **Table S2.** List of immune genes used in this study | Page 4 |
| **Table S3.** Population genetic statistics of immune genes in the North American population – all sites | Page 8 |
| **Table S4.** Population genetic statistics of immune genes in the Florida population – all sites | Page 9 |
| **Table S5.** Population genetic statistics of immune genes in the Pacific population – all sites | Page 10 |
| **Table S6.** Population genetic statistics of immune genes in the Atlantic population – all sites | Page 11 |
| **Table S7.** Immune genes are not disproportionally represented in genome-wide outliers | Page 12 |
| **Figure S1**. Population genetic statistics of immune genes in the North American population – all sites | Page 13 |
| **Figure S2.** Population genetic statistics of immune genes in all four populations – all sites | Page 14 |
| **Figure S3.** $F_{ST}$ of immune genes in each derived population compared to the ancestral population – all sites | Page 15 |

**Table S1.** *Danaus* samples used in this study. Sample information obtained from Zhan *et al.* (2014). We assigned each *D. plexippus* samples into genetic populations based on their collecting location.

| Species | Accession | Sample | Sex | Collecting location | Assigned population |
|---|---|---|---|---|---|
| *D. plexippus* | SRX681756 | Plex_CA_W48_F | female | Pismo Beach, CA, USA | North America |
| *D. plexippus* | SRX680103 | Plex_FLn_StM109_M | male | St. Marks, FL, USA | North America |
| *D. plexippus* | SRX680104 | Plex_FLn_StM122_F | female | St. Marks, FL, USA | North America |
| *D. plexippus* | SRX680105 | Plex_FLn_StM123_F | female | St. Marks, FL, USA | North America |
| *D. plexippus* | SRX681753 | Plex_FLn_StM146_M | male | St. Marks, FL, USA | North America |
| *D. plexippus* | SRX679271 | Plex_MA_HI012_F | female | Massachusetts, USA | North America |
| *D. plexippus* | SRX679303 | Plex_MA_HI023_M | male | Massachusetts, USA | North America |
| *D. plexippus* | SRX679310 | Plex_MA_HI035_F | female | Massachusetts, USA | North America |
| *D. plexippus* | SRX681737 | Plex_MEX_1742_F | female | Cerro Pelon, Mexico | North America |
| *D. plexippus* | SRX679305 | Plex_NJ_116_M | male | New Jersey, USA | North America |
| *D. plexippus* | SRX679306 | Plex_NJ_203_F | female | New Jersey, USA | North America |
| *D. plexippus* | SRX680106 | Plex_TX_T11_F | female | Texas, USA | North America |
| *D. plexippus* | SRX681744 | Plex_FLs_MIA2514_M | male | Miami, FL, USA | South Florida |
| *D. plexippus* | SRX681537 | Plex_FLs_MIA126_F | female | Miami, FL, USA | South Florida |
| *D. plexippus* | SRX681742 | Plex_FLs_MIA16_M | male | Miami, FL, USA | South Florida |
| *D. plexippus* | SRX681757 | Plex_FLs_MIA11_F | female | Miami, FL, USA | South Florida |
| *D. plexippus* | SRX681758 | Plex_FLs_MIA122_M | male | Miami, FL, USA | South Florida |
| *D. plexippus* | SRX681746 | Plex_FLs_MIA40_F | female | Miami, FL, USA | South Florida |
| *D. plexippus* | SRX681747 | Plex_FLs_MIA454_F | female | Miami, FL, USA | South Florida |
| *D. plexippus* | SRX680118 | Plex_WSM_M36_M | male | Samoa | Pacific |
| *D. plexippus* | SRX681528 | Plex_WSM_M38_F | female | Samoa | Pacific |
| *D. plexippus* | SRX681647 | Plex_FJI_M81_M | male | Fiji | Pacific |
| *D. plexippus* | SRX681648 | Plex_FJI_M82_M | male | Fiji | Pacific |
| *D. plexippus* | SRX681541 | Plex_NCL_M122_M | male | New Caledonia | Pacific |
| *D. plexippus* | SRX681543 | Plex_NCL_M128_M | male | New Caledonia | Pacific |
| *D. plexippus* | SRX681765 | Plex_AUS_18_F | female | Australia | Pacific |
| *D. plexippus* | SRX681766 | Plex_AUS_87_M | male | Australia | Pacific |
| *D. plexippus* | SRX681532 | Plex_AUS_99_F | female | Australia | Pacific |
| *D. plexippus* | SRX681767 | Plex_NZL_16_M | male | New Zealand | Pacific |
| *D. plexippus* | SRX681538 | Plex_NZL_3_F | female | New Zealand | Pacific |
| *D. plexippus* | SRX681768 | Plex_NZL_4_F | female | New Zealand | Pacific |
| *D. plexippus* | SRX680109 | Plex_ESP_28_M | male | Spain | Atlantic |
| *D. plexippus* | SRX680110 | Plex_ESP_29_F | female | Spain | Atlantic |
| *D. plexippus* | SRX680111 | Plex_ESP_30_M | male | Spain | Atlantic |
| *D. plexippus* | SRX681546 | Plex_MAR_M09_F | female | Morocco | Atlantic |
| *D. plexippus* | SRX681545 | Plex_MAR_M08_M | male | Morocco | Atlantic |
| *D. plexippus* | SRX681544 | Plex_MAR_M034_M | male | Morocco | Atlantic |
| *D. gilippus* | SRX998564 | Gili_TX_01_F | female | Texas, USA | N/A |

**Table S2.** List of immune genes used in this study. Note that immune genes on sex chromosomes were not included in the analyses.

| Gene ID | Gene name | Gene length | Functional class | Chromosome |
|---|---|---|---|---|
| DPOGS200905 | PGRP-like | 3371 | Recognition | Z-chromosome |
| DPOGS209813 | PGRP-like | 1574 | Recognition | Autosome |
| DPOGS206909 | PGRP-like | 1472 | Recognition | Z-chromosome |
| DPOGS206910 | PGRP-like | 7919 | Recognition | Z-chromosome |
| DPOGS207148 | PGRP-like | 15320 | Recognition | Z-chromosome |
| DPOGS209814 | PGRP-like | 2222 | Recognition | Autosome |
| DPOGS206026 | PGRP-like | 3333 | Recognition | Z-chromosome |
| DPOGS212963 | BGRP-like | 7393 | Recognition | Autosome |
| DPOGS215599 | BGRP-like | 2910 | Recognition | Autosome |
| DPOGS212940 | BGRP-like | 4002 | Recognition | Autosome |
| DPOGS212941 | BGRP-like | 3544 | Recognition | Autosome |
| DPOGS212964 | BGRP-like | 3508 | Recognition | Autosome |
| DPOGS212965 | BGRP-like | 3969 | Recognition | Autosome |
| DPOGS203317 | Frep-like | 5328 | Recognition | Z-chromosome |
| DPOGS206045 | Frep-like | 4504 | Recognition | Z-chromosome |
| DPOGS203951 | Frep-like | 10862 | Recognition | Z-chromosome |
| DPOGS210549 | Class B-like SCR | 5148 | Recognition | Autosome |
| DPOGS203180 | Class B-like SCR | 13541 | Recognition | Autosome |
| DPOGS202796 | Class B-like SCR | 9956 | Recognition | Autosome |
| DPOGS214397 | Other SCR | 13196 | Recognition | Autosome |
| DPOGS213636 | Other SCR | 15159 | Recognition | Autosome |
| DPOGS212634 | Other SCR | 14283 | Recognition | Autosome |
| DPOGS202826 | Other SCR | 9352 | Recognition | Autosome |
| DPOGS215836 | TEP-like | 10955 | Recognition | Autosome |
| DPOGS210251 | NIM-like | 13838 | Recognition | Autosome |
| DPOGS210210 | NIM-like | 5318 | Recognition | Autosome |
| DPOGS210211 | NIM-like | 13481 | Recognition | Autosome |
| DPOGS204835 | CLIP-like | 3824 | Modulation | Autosome |
| DPOGS205231 | CLIP-like | 27491 | Modulation | Autosome |
| DPOGS206561 | CLIP-like | 4655 | Modulation | Autosome |
| DPOGS215180 | CLIP-like | 11215 | Modulation | Autosome |
| DPOGS215181 | CLIP-like | 10878 | Modulation | Autosome |
| DPOGS206562 | CLIP-like | 14594 | Modulation | Autosome |
| DPOGS206563 | CLIP-like | 3009 | Modulation | Autosome |
| DPOGS213841 | CLIP-like | 3609 | Modulation | Autosome |
| DPOGS215183 | CLIP-like | 13397 | Modulation | Autosome |
| DPOGS215188 | CLIP-like | 4339 | Modulation | Autosome |

| | | | | |
|---|---|---|---|---|
| DPOGS204146 | CLIP-like | 5061 | Modulation | Autosome |
| DPOGS204147 | CLIP-like | 6895 | Modulation | Autosome |
| DPOGS201678 | CLIP-like | 3492 | Modulation | Autosome |
| DPOGS215220 | CLIP-like | 3818 | Modulation | Autosome |
| DPOGS215098 | CLIP-like | 27161 | Modulation | Autosome |
| DPOGS208169 | CLIP-like | 3911 | Modulation | Autosome |
| DPOGS201966 | CLIP-like | 10337 | Modulation | Autosome |
| DPOGS215182 | CLIP-like | 10004 | Modulation | Autosome |
| DPOGS211355 | CLIP-like | 3601 | Modulation | Autosome |
| DPOGS203664 | CLIP-like | 7166 | Modulation | Autosome |
| DPOGS210568 | CLIP-like | 8283 | Modulation | Autosome |
| DPOGS214570 | CLIP-like | 4377 | Modulation | Autosome |
| DPOGS204148 | CLIP-like | 2905 | Modulation | Autosome |
| DPOGS205210 | CLIP-like | 3342 | Modulation | Autosome |
| DPOGS211237 | CLIP-like | 2943 | Modulation | Autosome |
| DPOGS206224 | CLIP-like | 5720 | Modulation | Autosome |
| DPOGS206217 | CLIP-like | 3300 | Modulation | Autosome |
| DPOGS205206 | CLIP-like | 3310 | Modulation | Autosome |
| DPOGS209809 | SPZ-like | 1777 | Signaling - Toll | Autosome |
| DPOGS209810 | SPZ-like | 5018 | Signaling - Toll | Autosome |
| DPOGS203200 | Toll_like-receptors | 3821 | Signaling - Toll | Autosome |
| DPOGS205279 | Toll_like-receptors | 4664 | Signaling - Toll | Autosome |
| DPOGS202626 | Toll_like-receptors | 2709 | Signaling - Toll | Autosome |
| DPOGS205281 | Toll_like-receptors | 3894 | Signaling - Toll | Autosome |
| DPOGS205295 | Toll_like-receptors | 3887 | Signaling - Toll | Autosome |
| DPOGS205123 | Toll_like-receptors | 5897 | Signaling - Toll | Autosome |
| DPOGS211472 | Toll_like-receptors | 4533 | Signaling - Toll | Autosome |
| DPOGS200002 | Toll_like-receptors | 5428 | Signaling - Toll | Autosome |
| DPOGS205283 | Toll_like-receptors | 1949 | Signaling - Toll | Autosome |
| DPOGS215274 | Toll_like-receptors | 5667 | Signaling - Toll | Autosome |
| DPOGS203198 | Toll_like-receptors | 3410 | Signaling - Toll | Autosome |
| DPOGS205293 | Toll_like-receptors | 869 | Signaling - Toll | Autosome |
| DPOGS205296 | Toll_like-receptors | 2006 | Signaling - Toll | Autosome |
| DPOGS207788 | Tollip | 3828 | Signaling - Toll | Autosome |
| DPOGS205936 | MyD88 | 2965 | Signaling - Toll | Autosome |
| DPOGS208945 | Tube | 2121 | Signaling - Toll | Autosome |
| DPOGS214647 | Pellino | 3601 | Signaling - Toll | Autosome |
| DPOGS210260 | Pelle | 11577 | Signaling - Toll | Autosome |
| DPOGS202662 | TRAF2 | 6521 | Signaling - Toll | Autosome |
| DPOGS209243 | ECSIT | 1486 | Signaling - Toll | Autosome |

| | | | | |
|---|---|---|---|---|
| DPOGS209453 | Cactus | 2181 | Signaling - Toll | Autosome |
| DPOGS215778 | IMD | 1319 | Signaling - IMD | Autosome |
| DPOGS200403 | TAK1 | 12371 | Signaling - IMD | Autosome |
| DPOGS202907 | IKKgamma | 14063 | Signaling - IMD | Autosome |
| DPOGS202564 | IKKbeta | 1586 | Signaling - IMD | Autosome |
| DPOGS207960 | FADD | 916 | Signaling - IMD | Autosome |
| DPOGS212093 | Dredd | 1508 | Signaling - IMD | Autosome |
| DPOGS200977 | Tab2 | 3087 | Signaling - IMD | Autosome |
| DPOGS203759 | IAP2 | 5539 | Signaling - IMD | Autosome |
| DPOGS201405 | Ubc13 | 455 | Signaling - IMD | Autosome |
| DPOGS208954 | Hem | 3377 | Signaling - JNK | Autosome |
| DPOGS213169 | JNK | 3769 | Signaling - JNK | Autosome |
| DPOGS214573 | Fos | 2706 | Signaling - JNK | Autosome |
| DPOGS202887 | Jun | 242 | Signaling - JNK | Autosome |
| DPOGS214325 | PIAS | 12079 | Signaling - JAK-STAT | Autosome |
| DPOGS214451 | SOCS | 2617 | Signaling - JAK-STAT | Autosome |
| DPOGS200349 | DOMELESS | 2960 | Signaling - JAK-STAT | Autosome |
| DPOGS210157 | Hopscotch | 8831 | Signaling - JAK-STAT | Autosome |
| DPOGS212956 | Stat | 16032 | Signaling - JAK-STAT | Autosome |
| DPOGS213997 | Attacin-Like | 877 | Effector | Autosome |
| DPOGS205720 | Attacin-Like | 1439 | Effector | Autosome |
| DPOGS215451 | Attacin-Like | 818 | Effector | Autosome |
| DPOGS210270 | Cecropin-like | 374 | Effector | Autosome |
| DPOGS210268 | Cecropin-like | 422 | Effector | Autosome |
| DPOGS210269 | Cecropin-like | 529 | Effector | Autosome |
| DPOGS200256 | Cecropin-like | 428 | Effector | Autosome |
| DPOGS210271 | Cecropin-like | 352 | Effector | Autosome |
| DPOGS210265 | Cecropin-like | 848 | Effector | Autosome |
| DPOGS210304 | Gloverin-like | 644 | Effector | Autosome |
| DPOGS210303 | Gloverin-like | 822 | Effector | Autosome |
| DPOGS202093 | NOS-like | 15626 | Effector | Autosome |
| DPOGS202094 | NOS-like | 19723 | Effector | Autosome |
| DPOGS201818 | PPO-like | 4660 | Effector | Z-chromosome |
| DPOGS201819 | PPO-like | 5674 | Effector | Z-chromosome |
| DPOGS206820 | PPO-like | 9352 | Effector | Z-chromosome |
| DPOGS200017 | PPO-like | 5087 | Effector | Autosome |

**Table S3.** Population genetic statistics of immune genes in the North American population using the paired-control approach, based on all sites within each gene. The $F_{ST}$ section was non-applicable because the North American population was the reference population used for population comparisons. “All immune” indicates the full immune gene set. In each statistic, the first raw shows the test statistic of the immune gene group. The second row shows the proportion of 10,000 permutations in which the difference between the means of the immune gene group and the control set was positive. Percentages < 2.5% and > 97.5 % were labeled in bold. The third row shows the *P*-value. *P*-values < 0.05 were labeled in bold. Asterisks indicates: * < 0.05, ** < 0.01, *** < 0.001.

| | **All Immune** | **Recognition** | **Signaling** | **Modulation** | **Effector** |
|---|---|---|---|---|---|
| All sites | | | | | |
| π: test statistic | -0.05 | 0.03 | -0.17 | 0.05 | 0.03 |
| π: > 0 (%) | 22.49 | 90.47 | **0.00** | 94.55 | 84.97 |
| π: *P*-value. | 0.451 | 0.196 | **0.000*** ** | 0.109 | 0.295 |
| Watterson's θ: test statistic | -0.03 | 0.05 | -0.19 | 0.08 | 0.03 |
| Watterson's θ: > 0 (%) | 34.28 | 94.35 | **0.00** | **98.69** | 79.51 |
| Watterson's θ: *P*-value | 0.687 | 0.122 | **0.000*** ** | **0.028*** | 0.414 |
| Tajima's D: test statistic | -8.46 | 0.66 | -11.63 | -2.60 | 5.11 |
| Tajima's D: > 0 (%) | 4.09 | 63.89 | **0.01** | 12.66 | **98.27** |
| Tajima's D: *P*-value | 0.084 | 0.718 | **0.001**** | 0.250 | **0.030*** |
| $F_{ST}$: test statistic | NA | NA | NA | NA | NA |
| $F_{ST}$: > 0 (%) | NA | NA | NA | NA | NA |
| $F_{ST}$: *P*-value | NA | NA | NA | NA | NA |

**Table S4.** Population genetic statistics of immune genes in the Florida population using the paired-control approach, based on all sites within each gene. $F_{ST}$ was compared to the North American population. "All immune" indicates the full immune gene set. In each statistic, the first raw shows the test statistic of the immune gene group. The second row shows the proportion of 10,000 permutations in which the difference between the means of the immune gene group and the control set was positive. Percentages < 2.5% and > 97.5 % were labeled in bold. The third row shows the *P*-value. *P*-values < 0.05 were labeled in bold. Asterisks indicates: * < 0.05, ** < 0.01, *** < 0.001.

| | **All Immune** | **Recognition** | **Signaling** | **Modulation** | **Effector** |
|---|---|---|---|---|---|
| All sites | | | | | |
| π: test statistic | -0.07 | 0.04 | -0.17 | 0.05 | 0.01 |
| π: > 0 (%) | 13.75 | 92.24 | **0.00** | 94.41 | 59.58 |
| π: *P*-value. | 0.278 | 0.160 | **0.000*** ** | 0.111 | 0.844 |
| Watterson's θ: test statistic | -0.05 | 0.05 | -0.18 | 0.07 | 0.01 |
| Watterson's θ: > 0 (%) | 23.95 | 94.95 | **0.00** | **97.76** | 57.51 |
| Watterson's θ: *P*-value | 0.479 | 0.108 | **0.000*** ** | **0.046*** | 0.866 |
| Tajima's D: test statistic | -12.35 | 0.89 | -13.48 | -2.80 | 3.04 |
| Tajima's D: > 0 (%) | **0.99** | 67.38 | **0.02** | 9.40 | 87.62 |
| Tajima's D: *P*-value | **0.021*** | 0.650 | **0.000*** ** | 0.189 | 0.248 |
| $F_{ST}$: test statistic | -0.09 | 0.04 | -0.02 | -0.06 | -0.05 |
| $F_{ST}$: > 0 (%) | 35.52 | 75.72 | 48.40 | 34.08 | 25.14 |
| $F_{ST}$: *P*-value | 0.679 | 0.485 | 0.920 | 0.579 | 0.489 |

**Table S5.** Population genetic statistics of immune genes in the Pacific population using the paired-control approach, based on all sites within each gene. $F_{ST}$ was compared to the North American population. "All immune" indicates the full immune gene set. In each statistic, the first raw shows the test statistic of the immune gene group. The second row shows the proportion of 10,000 permutations in which the difference between the means of the immune gene group and the control set was positive. Percentages < 2.5% and > 97.5 % were labeled in bold. The third row shows the *P*-value. *P*-values < 0.05 were labeled in bold. Asterisks indicates: * < 0.05, ** < 0.01, *** < 0.001.

| | **All Immune** | **Recognition** | **Signaling** | **Modulation** | **Effector** |
|---|---|---|---|---|---|
| All sites | | | | | |
| π: test statistic | -0.03 | 0.04 | -0.14 | 0.04 | 0.04 |
| π: > 0 (%) | 29.57 | 97.12 | **0.00** | 92.75 | 86.69 |
| π: *P*-value. | 0.570 | 0.058 | **0.000*** ** | 0.139 | 0.243 |
| Watterson's θ: test statistic | -0.02 | 0.04 | -0.12 | 0.04 | 0.03 |
| Watterson's θ: > 0 (%) | 32.68 | **99.41** | **0.00** | 95.93 | 87.26 |
| Watterson's θ: *P*-value | 0.635 | **0.013*** | **0.000*** ** | 0.077 | 0.231 |
| Tajima's D: test statistic | -1.29 | -2.94 | 2.26 | -1.78 | 1.17 |
| Tajima's D: > 0 (%) | 43.76 | 16.54 | 68.12 | 32.22 | 62.68 |
| Tajima's D: *P*-value | 0.873 | 0.329 | 0.630 | 0.662 | 0.754 |
| $F_{ST}$: test statistic | -0.50 | -0.21 | 0.08 | -0.43 | 0.06 |
| $F_{ST}$: > 0 (%) | 26.09 | 24.37 | 58.39 | 16.57 | 58.58 |
| $F_{ST}$: *P*-value | 0.505 | 0.461 | 0.862 | 0.327 | 0.878 |

**Table S6.** Population genetic statistics of immune genes in the Atlantic population using the paired-control approach, based on all sites within each gene. $F_{ST}$ was compared to the North American population. “All immune” indicates the full immune gene set. In each statistic, the first raw shows the test statistic of the immune gene group. The second row shows the proportion of 10,000 permutations in which the difference between the means of the immune gene group and the control set was positive. Percentages < 2.5% and > 97.5 % were labeled in bold. The third row shows the *P*-value. *P*-values < 0.05 were labeled in bold. Asterisks indicates: * < 0.05, ** < 0.01, *** < 0.001.

| | **All Immune** | **Recognition** | **Signaling** | **Modulation** | **Effector** |
|---|---|---|---|---|---|
| All sites | | | | | |
| π: test statistic | -0.04 | 0.00 | -0.12 | 0.03 | 0.05 |
| π: > 0 (%) | 24.54 | 58.07 | **0.00** | 83.25 | 92.68 |
| π: *P*-value. | 0.483 | 0.852 | **0.001**** | 0.324 | 0.103 |
| Watterson's θ: test statistic | 0.00 | 0.01 | -0.10 | 0.04 | 0.04 |
| Watterson's θ: > 0 (%) | 47.51 | 78.50 | **0.01** | 93.64 | 93.63 |
| Watterson's θ: *P*-value | 0.940 | 0.447 | **0.001**** | 0.122 | 0.087 |
| Tajima's D: test statistic | -21.03 | -6.60 | -8.49 | -6.55 | 0.61 |
| Tajima's D: > 0 (%) | **1.53** | 6.75 | 8.62 | 8.51 | 55.26 |
| Tajima's D: *P*-value | **0.029*** | 0.119 | 0.167 | 0.159 | 0.880 |
| $F_{ST}$: test statistic | 1.16 | 0.30 | 1.37 | 0.07 | -0.57 |
| $F_{ST}$: > 0 (%) | 91.75 | 81.55 | **99.38** | 57.80 | 3.48 |
| $F_{ST}$: *P*-value | 0.156 | 0.368 | **0.009**** | 0.875 | 0.094 |

**Table S7.** Immune genes are not disproportionally represented in genome-wide outliers. We used chi-square tests to evaluate whether immune genes (n = 102) are disproportionally represented in genome-wide outliers in each population at either 0-fold or 4-fold degeneracy sites. Outliers were defined as < 2.5$^{th}$ percentile or > 97.5$^{th}$ percentile of the genome background in either Tajima's D or $F_{ST}$. For the North American population, outliers were identified based on only the Tajima's D data.

| **Population** | **Sites** | **Number of outlier immune genes** | **$\chi$2** | ***df*** | **P-value** |
|---|---|---|---|---|---|
| North America | 0-fold | 3 | 0.47 | 1 | 0.49 |
| | 4-fold | 5 | 0.00 | 1 | 1.00 |
| South Florida | 0-fold | 13 | 0.86 | 1 | 0.35 |
| | 4-fold | 9 | 0.00 | 1 | 1.00 |
| Pacific | 0-fold | 8 | 0.16 | 1 | 0.69 |
| | 4-fold | 4 | 2.67 | 1 | 0.10 |
| Atlantic | 0-fold | 9 | 0.01 | 1 | 0.91 |
| | 4-fold | 13 | 0.59 | 1 | 0.44 |

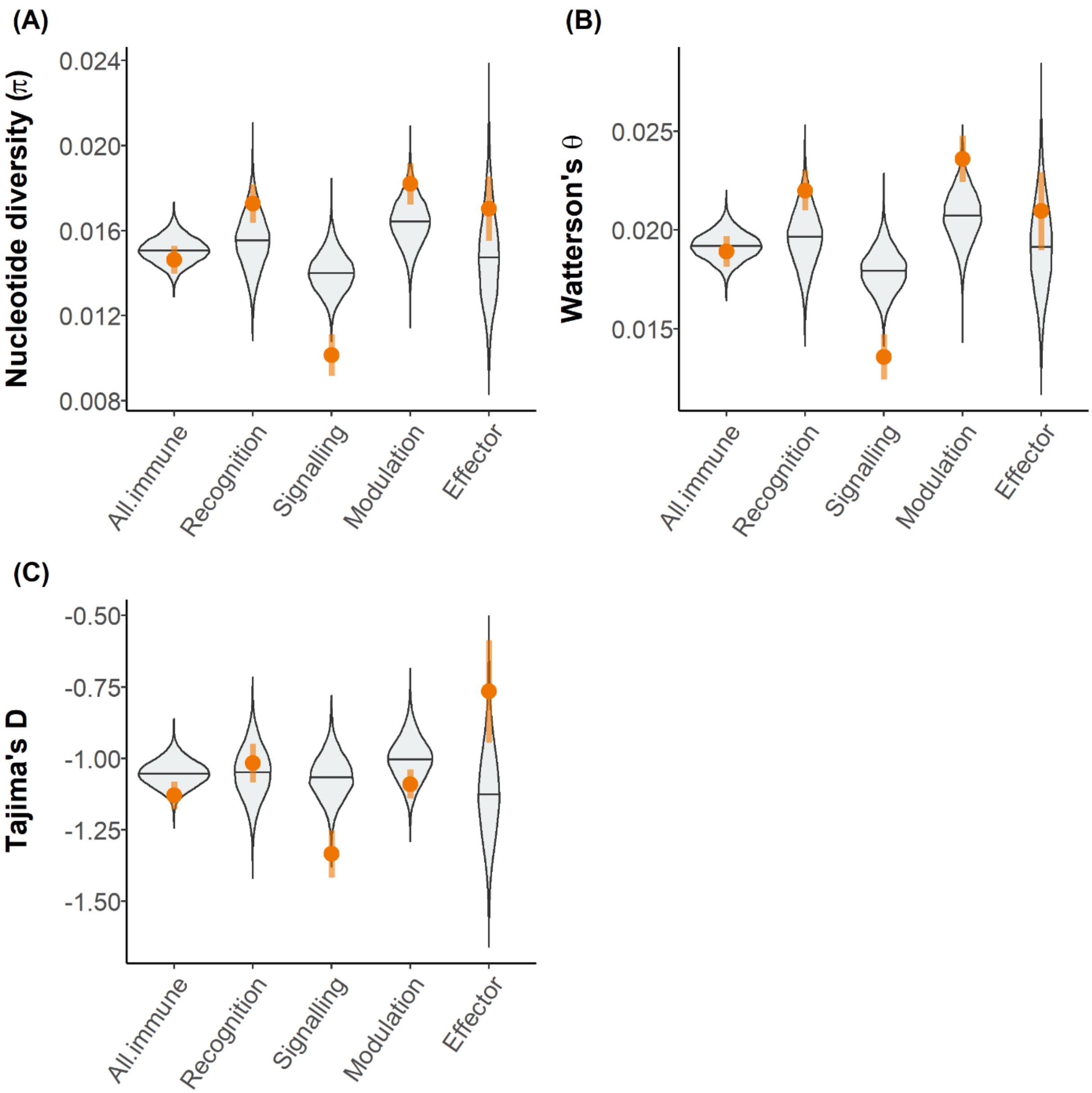


**Figure S1.** Population genetic statistics of immune genes in the North American population using the paired-control approach, based on all sites within each gene. (a): Nucleotide diversity (**π**); (b): Watterson's θ; (c): Tajima's D. Each immune gene group was compared to selected pair-control sets. Violin plots show the distribution of the mean of each control set generated with 10, 000 permutations. The orange dots and vertical lines indicate mean ±1 SEM of the immune gene group of interest.

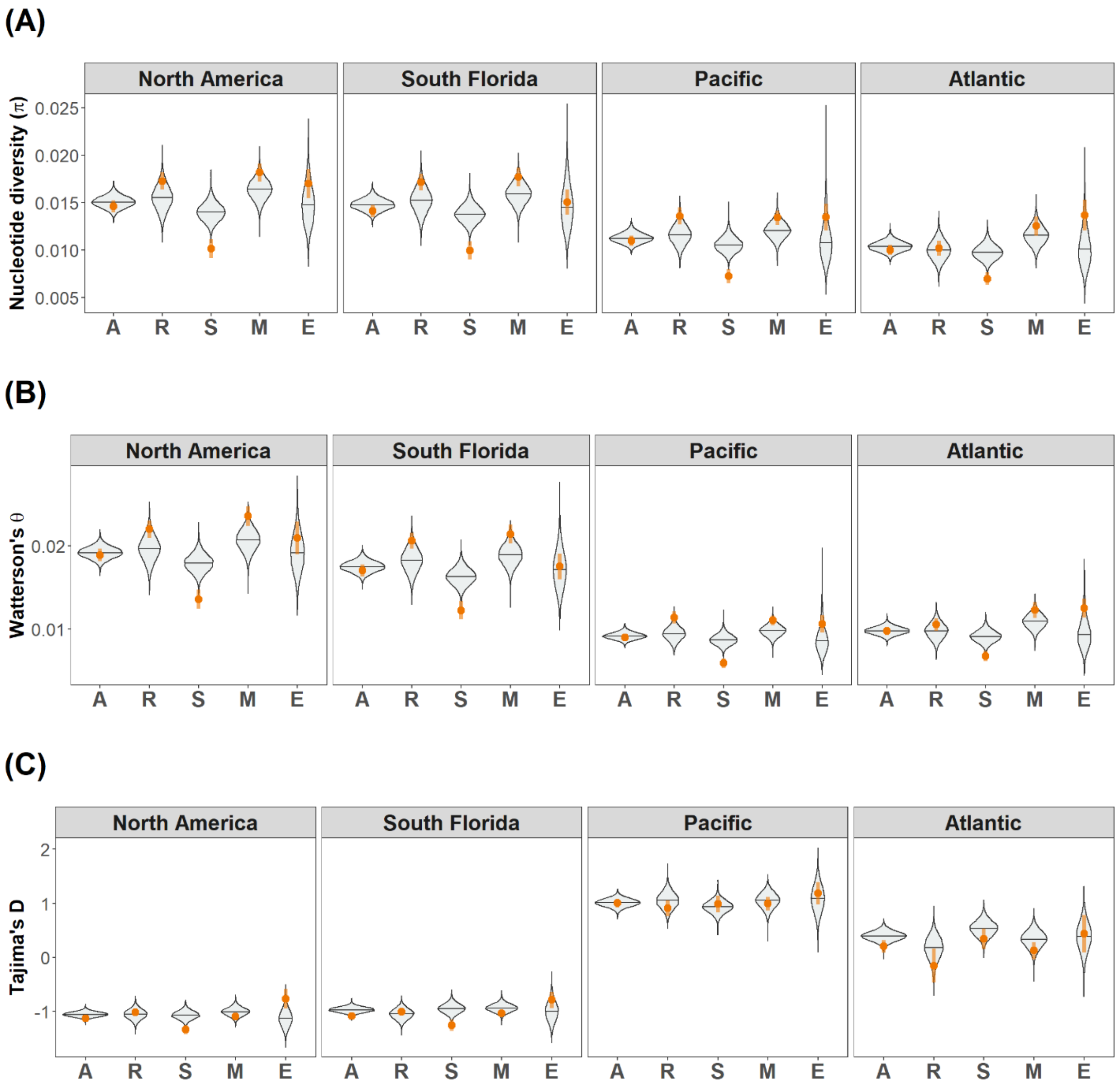


**Figure S2.** Population genetic statistics of immune genes in all four populations (North America, Florida, Pacific, and Atlantic) using the paired-control approach, based on all sites within each gene. (a): Nucleotide diversity (**$\pi$**); (b): Watterson's θ; (c): Tajima's D. Each immune gene group was compared to selected pair-control sets. Violin plots show the distribution of the mean of each control set generated with 10, 000 permutations. The orange dots and vertical lines indicate mean ±1 SEM of the immune gene group of interest. X-axis represents immune gene groups: all immune genes (A), recognition genes (R), signaling genes (S), modulation genes (M), and effector genes (E).

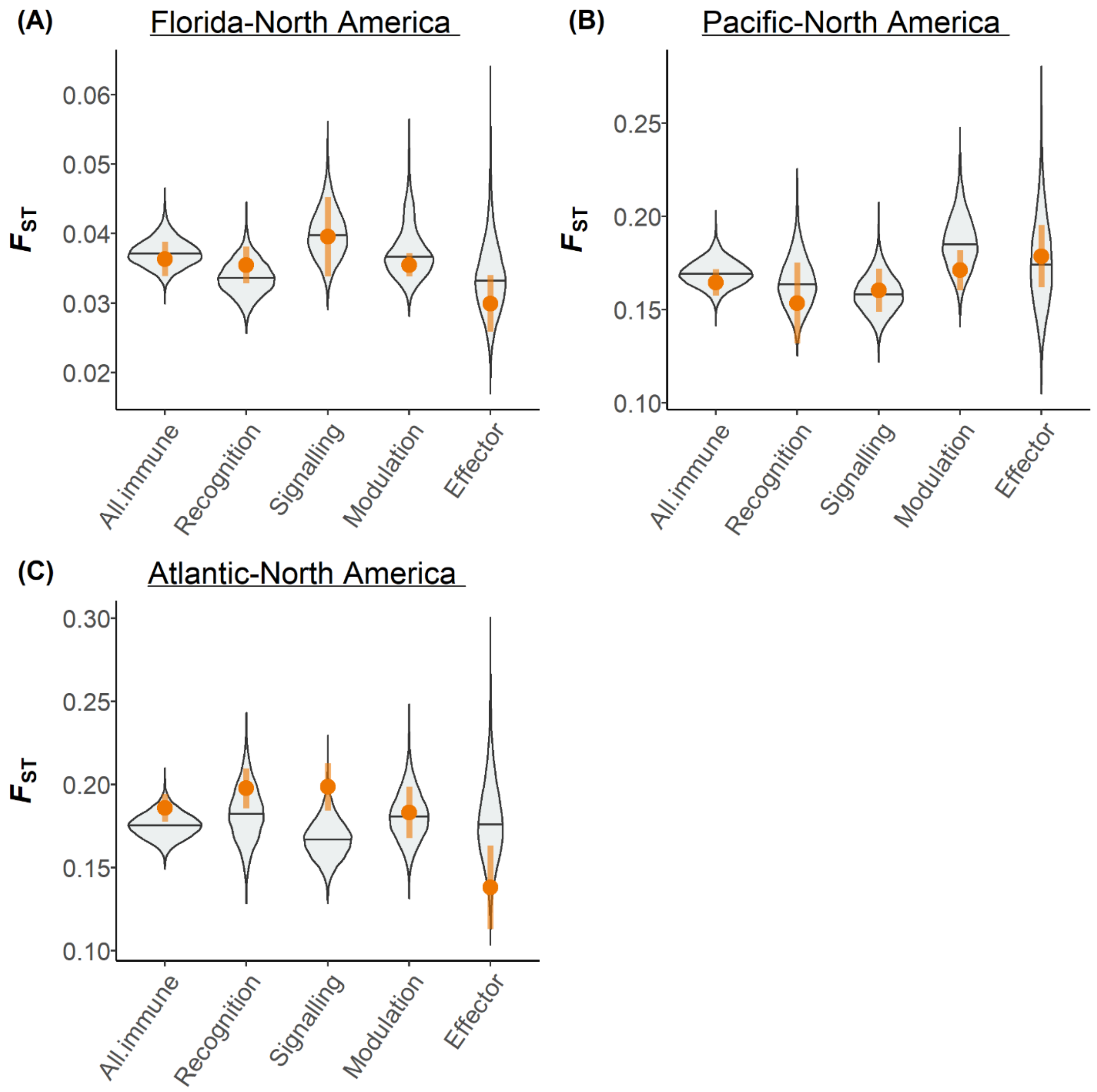


**Figure S3.** $F_{ST}$ of immune genes in each derived population compared to the ancestral (North American) population using the paired-control approach, based on all sites within each gene. (a): South Florida population (**$\pi$**); (b): Pacific population; (c): Atlantic population. Each immune gene group was compared to selected pair-control sets. Violin plots show the distribution of the mean of each control set generated with 10, 000 permutations. The orange dots and vertical lines indicate mean ±1 SEM of the immune gene group of interest.
